# The genomic view of diversification

**DOI:** 10.1101/413427

**Authors:** Julie Marin, Guillaume Achaz, Anton Crombach, Amaury Lambert

## Abstract

Evolutionary relationships between species are traditionally represented in the form of a tree, the species tree. Its reconstruction from molecular data is hindered by frequent conflicts between gene genealogies. Usually, these disagreements are explained by incomplete lineage sorting (ILS) due to random coalescences of gene lineages inside the edges of the species tree. This paradigm, the multi-species coalescent (MSC), is constantly violated by the ubiquitous presence of gene flow, leading to incongruences between gene trees that cannot be explained by ILS alone. Here we argue instead in favor of a vision acknowledging the importance of gene flow and where gene histories shape the species tree rather than the opposite. We propose a new framework for modeling the joint evolution of gene and species lineages relaxing the hierarchy between the species tree and gene trees. We implement this framework in two mathematical models called the gene-based diversification models (GBD): 1) GBD-forward following all evolving genomes and 2) GBD-backward based on coalescent theory. They feature four parameters tuning colonization, gene flow, genetic drift and genetic differentiation. We propose a quick inference method based on differences between gene trees. Applied to two empirical data-sets prone to gene flow, we find a better support for the GBD model than for the MSC model. Along with the increasing awareness of the extent of gene flow, this work shows the importance of considering the richer signal contained in genomic histories, rather than in the mere species tree, to better apprehend the complex evolutionary history of species.

## Introduction

The most widely used way of representing evolutionary relationships between contemporary species is the so-called species tree, or phylogeny. The high efficiency of statistical methods using sequence data to reconstruct species trees, hence called ‘molecular phylogenies’, led to precise dating of the nodes of these phylogenies (Heled & Drummond, 2010; Kishino, Thorne, & Bruno, 2001; Tamura et al., 2012). Notwithstanding the debatable accuracy of these datings, the use of time-calibrated phylogenies, sometimes called ‘timetrees’ (Hedges & Kumar, 2009), has progressively overtaken a view where phylogenies merely represent tree-like relationships between species in favor of a view where the timetree is the exact reflection of the diversification process (Morlon, 2014; Pyron & Burbrink, 2013; Stadler, 2013a). In this view, the nodes of the phylogeny are consequently seen as punctual speciation events where one daughter species is instantaneously ‘born’ from a mother species. In this paper, we explore an alternative view of diversification, acknowledging that speciation is a long-term process (Etienne, Morlon, & Lambert, 2014; Lambert, Morlon, & Etienne, 2015; Rosindell, Cornell, Hubbell, & Etienne, 2010) and not invoking any notion of mother-daughter relationship between species as done in the timetree view. This alternative view is gene-based rather than species-based, comparable with Wu’s genic view of speciation (2001). We use here the term ‘gene’ in the sense of “non-recombining locus”, *i.e.*, a region of the genome with a unique evolutionary history. Our view is meant in particular to accommodate the well-recognized existence of gene flow between incipient species, which persists during the speciation process and long after (Mallet, Besansky, & Hahn, 2016).

The timetree view of phylogenies does acknowledge that gene trees are not independent and may disagree with the species tree (Maddison, 1997). However, current methods jointly inferring gene trees and species tree rely on two assumptions that we question in the next section: (1) there is a unique species tree, (2) the species tree shapes the gene trees and (3) the species tree is the only factor mediating all dependencies between gene trees (they are independent conditional on the species tree).

This view is materialized in a model called the ‘multispecies coalescent’ (MSC) (Knowles & Kubatko, 2011) where conditional on the species tree, the evolutionary histories of genes follow independent coalescents constrained to take place within the hollow edges of the species tree. Many methods have been developed to estimate the species tree under the MSC, such as full likelihood methods (e.g. beast, Heled and Drummond 2010, bpp, Yang 2015) which average over gene trees and parameters (Xu & Yang, 2016), and the approximate or summary coalescent methods (e.g. Astral, Mirarab et al. 2014, mp-est, L. Liu, Yu, and Edwards 2010, and Stells, Y. Wu 2012) which use a two-step approach: gene trees are first inferred and then combined to estimate the species tree that minimize conflicts among gene trees. Discordance between gene topologies is then explained, as a first approximation at least, by the intrinsic randomness of coalescences resulting in incomplete lineage sorting (ILS) (Fig. 1).

**Figure 1:**
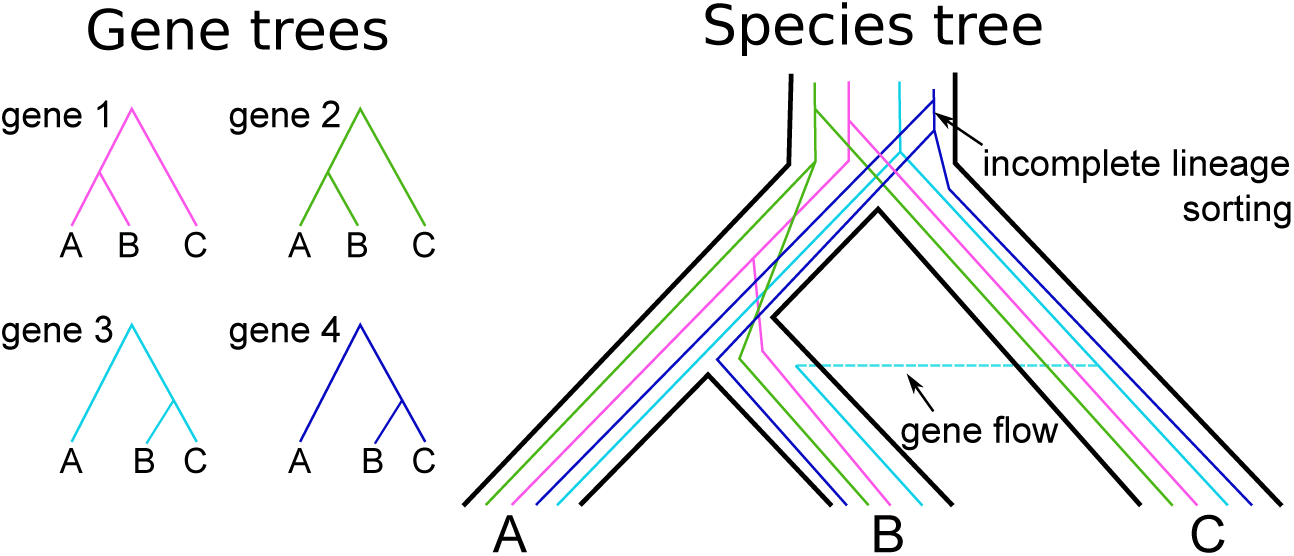
Gene trees and species tree conflicts. The species tree of A, B, and C is depicted in black. In pink (gene 1) and green (gene 2) are two gene trees congruent with the species tree, *i.e.* with A and B being sister species. In light blue (gene 3), the tree of a gene subject to gene flow between species B and C. In dark blue (gene 4), the tree of a gene undergoing incomplete lineage sorting.

However, the presence of gene flow (introgression, hybridization, horizontal transfer) is now widely recognized between closely related species, and even between distantly related species (Mallet et al., 2016). Porous species boundaries, allowing for gene exchange because of incomplete reproductive isolation, are indeed regularly observed in diverse taxa such as amphibians (Fontenot, Makowsky, & Chippindale, 2011; Pereira, Monahan, & Wake, 2011), arthropods (De Busschere et al., 2010), cichlids (Willis, Macrander, Farias, & Ortí, 2012), cyprinids (Buonerba et al., 2015; Gante, Collares-Pereira, & Coelho, 2004; Gante, Doadrio, Alves, & Dowling, 2015; Gante, Santos, & Alves, 2010; Sousa-Santos et al., 2014), insects (Nadeau et al., 2013; Peccoud, Ollivier, Plantegenest, & Simon, 2009; Wahlberg, Weingartner, Warren, & Nylin, 2009), and even more frequently among bacteria (Mallet et al., 2016; Soucy, Huang, & Gogarten, 2015). Long neglected, gene flow has recently been recognized as an important evolutionary driving force, through adaptive introgression or the formation of new hybrid taxa (Abbott et al., 2013). The ubiquity of genetic exchange across the Tree of Life between contemporary species suggests that gene flow has occurred many times in the evolutionary past, and might actually be the most important cause of discrepancies between gene histories (e.g. Clark and Messer 2015; Cui et al. 2013; Gallus, Janke, Kumar, and Nilsson 2015; Jónsson et al. 2014) (Fig. 1). Accordingly, several extensions to the MSC model have been considered allowing for gene flow between species (Kubatko, 2009; Yu, Dong, Liu, & Nakhleh, 2014). These models acknowledge that species boundaries can be permeable at a few specific timepoints (Harrison & Larson, 2014). Unfortunately, because of the heavy computational cost of modeling the coalescent with gene flow, these methods are limited to small data-sets (Yu et al., 2014). More importantly, they might not be appropriate to realistically model gene flow, given the frequency of gene flow across time and clades described in empirical studies (Solís-Lemus, Yang, & Ané, 2016). Additionally, some of these methods, for instance astral and mp-est, might infer erroneous gene trees when gene flow is present (Long & Kubatko, 2018). These observations urge for novel approaches where gene flow is the rule rather than the exception.

To fill this void, we propose here an alternative framework and two accompanying models (one in forward time and one in backward time), the gene-based diversification (GBD) models, framed with minimal assumptions arising from recent empirical evidence. Those models rely on the property of populations to spontaneously differentiate genetically while simultaneously undergoing gene flow. This genetic differentiation is accompanied by a decrease in gene flow until reproductive isolation is complete. Moreover, unlike previous models, we place ourselves in the case of pervasive gene flow among species that may have occurred countless times in the past, as suggested by recent studies. The GBD models are anchored in a new conceptual framework, that we call the genomic view of diversification. Unlike the timetree view, the present framework does not put the emphasis on the species tree (which in our model becomes a network rather than a tree) and assumes that gene trees shape the species tree rather than the opposite.

## The genomic view of diversification

### Gene flow and the questionable existence of a species genealogy

The biological species concept (BSC, Mayr 1942) defines species as groups of interbreeding populations that are reproductively isolated from other groups. This definition postulates the non-permeability of species boundaries, which is contradicted by the growing body of evidence describing permeable or semi-permeable genomes, even between distantly related taxa. To integrate the possibility of gene flow into the definition of species, Wu (2001) shifted the emphasis from isolation at the level of the whole genome to differential isolation at the gene level. Species are thus defined as differentially adapted groups for which inter-specific gene flow is allowed except for genes involved in differential adaptation (a well-defined form of divergence in which the alternative alleles have opposite fitness effects in the two groups). Because a fraction of the genome may still be exchanged after speciation is complete, a mosaic of gene genealogies is expected between divergent genomes (Wu, 2001). Much evidence supports this prediction with the observation of highly conflicting gene trees, e.g. Darwin’s finches (B. R. Grant & Grant, 1998; P. R. Grant & Grant, 1996), sympatric sticklebacks (Rundle, Nagel, Boughman, & Schluter, 2000; Schluter, 1998), Iberian barbels (Gante et al., 2015), and *Rhagoletis* species (Berlocher, 2000).

Accordingly, the notion of a species genealogy as the binary division of species into new independently evolving lineages in bifurcating phylogenetic trees, appears inappropriate. To avoid this misleading vision of speciation, we here wish to relax the species tree constraint by considering only gene genealogies as real genealogies, thereby laying aside, at least temporarily, the notion of species genealogy. To do so, we do not specify mother-daughter relationships between species, yet we postulate the existence of species at any time, and assume that we can unambiguously follow the genealogies of genes (defined as non-recombining loci, as mentioned above).

The notion of a species genealogy as a binary bifurcating tree is hardly compatible with gene flow, and a direct consequence is to challenge the notion of a unique ancestral species. If all genes ancestral to species *S* have travelled through the same species in the past, then species *S* has only one single ancestor species at any time. But because of gene flow, these genes may lie in different species living at a given time in the past, such that species *S* can have several ancestral species at this time. In other words, several species have contributed to the present-day genome of the species *S*.

### Genomic coadaptation under continuous gene flow

While some genes (e.g., genes involved in divergent adaptation) are hardly exchanged between populations, other genes (e.g., neutral genes unlinked to genes under divergent selection) can be subject to gene flow between different species (Pinho & Hey, 2010; Wu, 2001). Gene flow can persist for long periods of time, with evidence suggesting introgression events occurring over periods lasting up to 20 Myr (Buonerba et al., 2015; Gante et al., 2015; Willis et al., 2012). Over time, genetic differences will accumulate in regions of low recombination and expand via selective sweeps, leading eventually to complete reproductive isolation (Wu, 2001). Because populations differentially accumulate new alleles, their compatibility (hybrid fitness) will be affected. This process has been conceptualized by Bateson, Dobzhanzky and Muller in the so-called Bateson-Dobzhanzky-Muller (BDM) model (Coyne & Orr, 2004; Dobzhansky, 1936; Muller, 1942). This model proposes that genetic incompatibilities, hence called BDM incompatibilities, are characterised by negative epistatic interactions between alleles at two or more genes that have fixed differentially, in each of the parental populations, by local adaptation or genetic drift. The selective value of hybrids is reduced because the new alleles, divergently selected in each population, are incompatible when carried by the same genome. On the other hand, in the parental populations the co-adapted combinations of alleles have neutral or even beneficial effects (Seehausen et al., 2014; Turelli & Orr, 2000). These incompatibilities have been hypothesized to increase at a rate proportional to the square of time (Orr, 1995). Accordingly, pairs of species will likely exhibit greater genetic incompatibility as a function of time since divergence, *i.e.* be less permeable to gene flow, as has been observed for Iberian barbels (Gante et al., 2015), pea aphids (Peccoud et al., 2009), or salamanders (Pereira et al., 2011). In other words, gene lineages remaining too long isolated within different species decrease their ability to introgress the genome of the other, a property that we name *genomic coadaptation* and which is the consequence of spontaneous mutation.

### The gene-based diversification (GBD) models

We propose here a new plastic framework, derived from the genomic view of diversification described above, that acknowledges the importance of gene flow and relaxes the hierarchy between the species tree and gene trees. We built two models, one in forward time that follows the standard view of the main biological processes responsible for diversification under gene flow, and one in backward time using coalescent theory, less computationally intensive, with matching backward parameters (Fig. 2). These models that we named the gene-based diversification (GBD-forward and GBD-backward) models, describe the joint evolution of gene and species lineages, reconciling phylogenomics with our current knowledge of species diversification. The biological mechanisms first, then the corresponding parameters, are detailed thereafter for each model.

**Figure 2:**
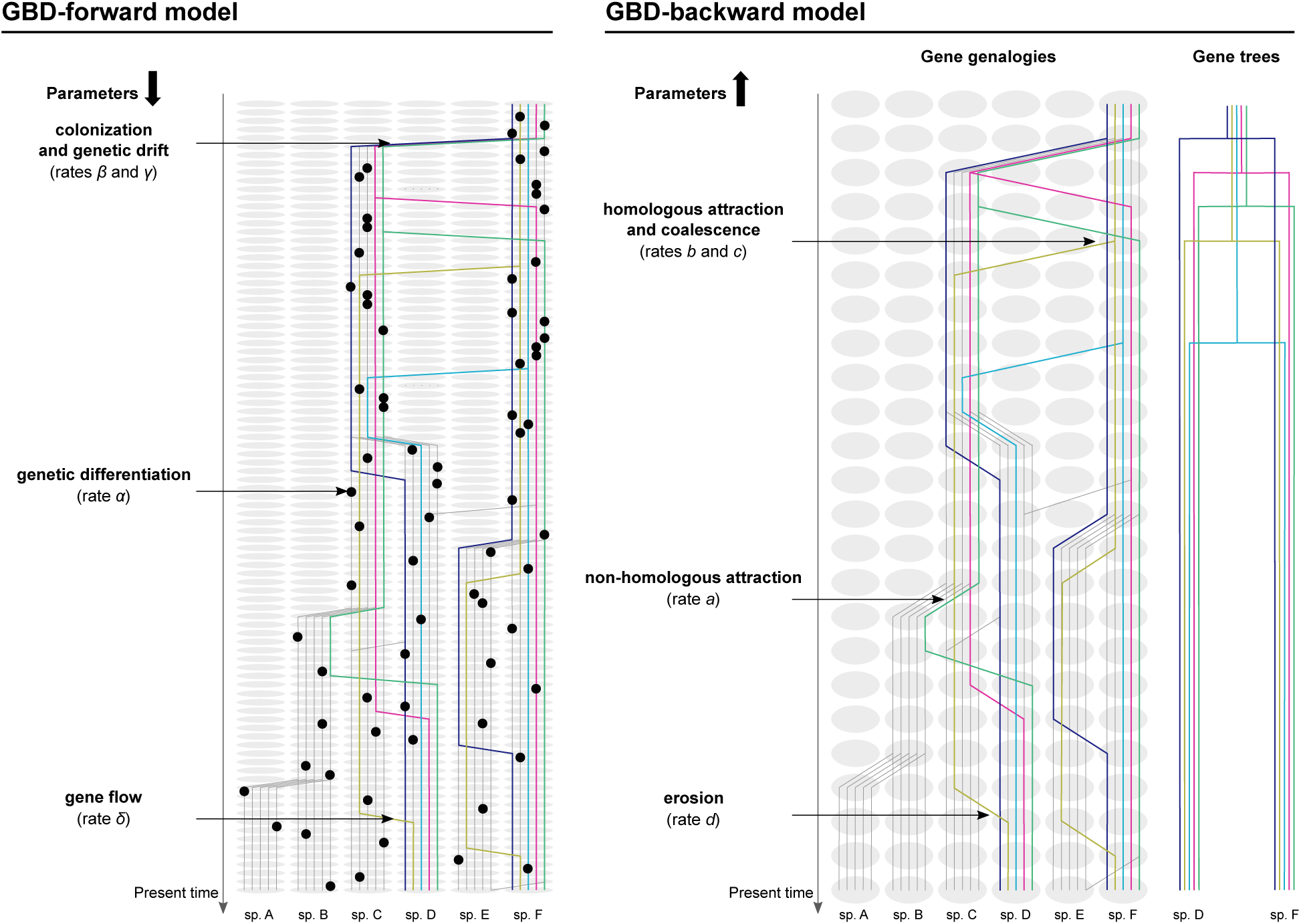
The gene-based diversification (GBD) models. Gene genealogies through species (or populations, depending on the point of view, retrospective *vs* prospective) are depicted for two hypothetical present-day genomes (*N* = 2 at *t* = 0) and five homologous genes (*n* = 5). Each grey ellipse represents a species (A-F). The grey lines represent the gene genealogies of non-sampled species at *t* = 0. The model assumes that species are quasi-static in the timescale of a few generations, and each species lineage is located in a separate column. The genealogies of genes depend on four processes: genetic differentiation/non-homologous attraction, colonization/homologous attraction, genetic drift/coalescence and gene flow/erosion.

### The GBD-forward model

The GBD-forward model describes the joint action of four processes affecting the diversification of genomes (see Fig. 2): colonization, mutation, drift and gene flow.

We consider a stochastically varying number of *populations*, all populated with individual genomes. We neglect extinctions and focus on *colonization* events, at which one population seeds a daughter population founded by one or several of its individuals. Genes independently accumulate *mutations* with time, under the infinite-allele model assumption. Mutations can be fixed or lost due to selection and genetic drift, that we summarize here under the term *drift*.

As a result of mutations and drift, populations differentiate genetically through time, which results in the decrease of gene flow. To model this, we follow what we term the *co-adaptation* between non-homologous genes and assume that *introgression* is governed by the numbers of co-adapted alleles in the receiver and donor populations. Right after colonization, all the genes of the daughter and mother populations carry the same alleles and so are co-adapted. Now an allele having arisen at time *t* by mutation on some gene is co-adapted only with the alleles carried by its genome at time *t*. This assumption underlies the well-known model of BDM incompatibilities described previously. Each time a mutation occurs the number of co-adapted genes among populations will decrease, reducing in turn the possibility of genetic exchange between populations.

Two populations that are completely differentiated, in the sense that all pairs of non-homologous alleles sampled from each of them are not co-adapted, can no longer exchange genes and can thus be seen as different species. Because populations are constantly differentiating from each other, we name populations in the prospective point of view (GBD-forward) what will become species only from a retrospective point of view (GBD-backward).

Demographic events are assumed to be much faster than other processes. In the time scale considered here, (1) the fixation of alleles within populations is instantaneous so that all genomes in a population are identical (we thus do not model the co-existence of several different homologous alleles within a population) and (2) a colonization event can be seen as the instantaneous replication of one population into two, actually because of (1), of one genome into two.

#### Parametrization

At *t* = 0, we consider a single monomorphic population, summarized into a single genome harboring *n* genes. During the diversification process, the genome of this population (*n* genes) will be replicated, mutations will be differentially fixed in each population, and the genomes of these populations can be replicated again. We follow the lineages of these *n* genes in forward time, assuming a time-continuous Markov chain with 4 events occurring at the following rates.

- **Genetic differentiation** (rate *α*). At any time *t*, each gene lineage in each population can acquire a new allele (infinite-allele model) at rate *α*. By definition, a new allele occurring at gene *L* on genome *G* is co-adapted with the allele present at a gene *L′*, for any *L′* (different of *L*) of genome *G*. On the contrary, a mutation arising at gene *L* of genome *G* and a mutation arising at gene *L′* of genome *G′* are not co-adapted.
- **Colonization** (rate *β*). At any time *t*, each population can be replicated at rate *β* into a new population which will evolve independently in the future. The newborn population is assumed to carry the same genome as carried by the mother population.
- **Genetic drift** (rate *γ*). Each population undergoes Moran-type births and deaths at rate *γ*. In this work, we assume *γ* to be much larger than all other parameters, so that each population is actually monomorphic at all times.
- **Gene flow** (rate *δ*). At any time *t*, each gene lineage at locus *L* on genome *G* can be replicated and introgress genome *G′* at rate *δ*(*n −* 1), proportional to the number of non-homologous loci in genome *G′*. If accepted by the target genome *G′*, the replicated lineage replaces its homologous gene lineage (at locus *L* in *G′*). The introgression is accepted with a probability equal to the fraction of the *n −* 1 non-homologous genes on *G′* carrying an allele co-adapted with the allele carried by *L*.

Diversification occurs until a number *K* of different populations is reached and the whole process is stopped when the *K* populations are genetically isolated, that is, when no pair of alleles carried by different genomes is co-adapted (i.e., when all probabilities of introgression are equal to 0).

This framework can be made more complex by letting the parameters depend on time, on the gene, or on any prescribed category of genes.

##### The GBD-backward model

The GBD-backward model is not the exact backward picture of the GBD-forward model but relies on the same idea that genomes in different populations tend to diverge with time until they cannot exchange alleles. The consequence of this fact is that genes sampled in the same genome today will tend to be found in the same population in the past more often than by chance. We model this phenomenon by saying that the ancestral lineages of genes sampled in the same present-day genome are *co-adapted*, and that co-adapted genes are *attracted* towards each other. The GBD-backward model describes the joint action of four processes (see Fig. 2): non-homologous attraction, homologous attraction, coalescence and erosion.

As explained above, in the retrospective point of view (GBD-backward), we name species the populations in which the ancestral gene lineages travel.

Each gene lineage can move from its species to another species. This happens as a result of homologous attraction, non-homologous attraction and erosion. As explained previously, (non-homologous) *co-adapted* genes move into the same species as a result of *non-homologous attraction*, which can be viewed as the backward consequence of *reproductive isolation*. Homologous gene lineages move into the same species as a result of *homologous attraction*, which can be viewed as the backward picture of a *colonization* event, when populations and their genomes have been replicated. Last, any gene lineage can move from its species by *erosion* to an empty species, i.e., a species containing no other gene lineage ancestral to the sample (the term erosion refers to the fact that the block of ancestral lineages lying in the same species loses one element).

When two homologous gene lineages are in the same species they can *coalesce* when finding their common ancestor, that is merge into a single lineage (hence within the same genome).

Note that after coalescence of two homologous lineages, the resulting lineage is now ancestral to at least two genomes and thus co-adapted with all gene lineages ancestral to these genomes. As a consequence of the mere *non-homologous attraction*, going further back in time, all other genes will then move to the same species and further coalesce, until all homologous gene lineages have coalesced.

Equivalently to the *drift* process in forward time, we will assume that *coalescences* are fast, so that in backward time homologous attraction events are immediately followed by coalescence of the two gene lineages.

#### Parameterization

At *t* = 0, *n* homologous genes are sampled in each of *N* distinct species. Retrospectively, the genomes of these *N* species (each harbouring *n* genes) will merge progressively into one genome of *n* genes at some time *t* in the past. Homologous genes, one by one, will merge (*homologous attraction* and *coalescence*). Merged genes will then attract all the genes of their original genomes (*non-homologous attraction*), until the *coalescence* of all homologous genes. We follow the lineages of these *n* genes in backward time, assuming a time-continuous Markov chain with 4 events occurring at the following rates.

- **Non-homologous attraction** (rate *a*). At any time *t* in the past, as a consequence of **genomic coadaptation**, each gene lineage *L* escapes from its species *S* at rate *a*(*n −* 1) per target species *S′*, proportional to the number of non-homologous loci in the genome *G′* hosted by *S′*. It is accepted in *S′* based on its co-adaptation with *G′*. If *G*_0_ denotes the genome harboring the descendant lineage of *L* at time *t* = 0, then all gene lineages harbored by *G′* that are ancestral to *G*_0_ are said co-adapted with *L*. Then *L* is accepted in *S′* with a probability proportional to the fraction of the *n −* 1 non-homologous loci of *G′* that are co-adapted with it. The parameter *a* corresponds to the parameter *α* of the GBD-forward model (genetic differentiation).
- **Homologous attraction** (rate *b*). At any time *t* in the past, each gene lineage at rate *b* per homologous gene lineage, moves to the species harboring this homologous lineage (or in an alternative, more specific version of the model, each gene lineage belonging to some previously prescribed category, like genes contributing to reproductive isolation). This parameter corresponds to the parameter *β* of the GBD-forward model (colonization).
- **Coalescence** (rate *c*). At any time *t* in the past, each pair of homologous genes lying within the same species coalesces at rate *c*. This parameter corresponds to the parameter *γ* of the GBD-forward model (genetic drift).
- **Erosion** (rate *d*). At any time *t* in the past, each gene lineage escapes from its genome at rate *d* and enters an empty species (also called ghost species, *i.e.*, harboring no other gene lineage ancestral to the samples, see Fig. 2). This parameter corresponds to the parameter *δ* of the GBD-forward model (gene flow). To model the flow of bigger chunks of DNA, we could alternatively assume that instead of one lineage, a given fraction of the lineages of a genome can simultaneously move to an otherwise empty species. We will not consider this possibility in the present work.

We define the number of *ancestral species* of a given genome at time *t*, as the number of species at time *t* containing gene lineages ancestral to this genome. Let us briefly expose the expected effects of these parameters on gene trees.

- Large non-homologous attraction (*a*) values will result in most gene lineages concentrated in one or two ancestral species.
- Large homologous attraction (*b*) values will result in short waits between speciation events.
- Large coalescence (*c*) values hinder incomplete lineage sorting.
- Large erosion (*d*) values will result in a large number of ancestral species per genome.

In this manuscript we wish to explore the impact of gene flow rather than ILS to explain gene tree conflicts, and thus consider a large *c* value (coalescence rate) so that coalescence events are instantaneous, which is consistent with the large *γ* value of the forward model. Therefore, only the parameters *a*, *b*, and *d* have an influence on the gene genealogies in the GBD-backward model.

The GBD models were implemented in R (https://www.r-project.org) and evaluated under different sets of parameters. Because the GBD-forward model is computationally prohibitive, while giving comparable qualitative results with the GBD-backward model (see Results), we conducted most of the analyses and the inferences with the GBD-backward model. We provide a first ABC-like inference method by minimizing the difference (Kullback-Leibler divergence) between the distributions of Kendall and Colijn (KC) distances (pairwise distances between gene trees) (Kendall & Colijn, 2016) in empirical vs simulated data. We applied this inference method to two empirical multi-locus data-sets showing complex evolutionary patterns due to gene flow, comprising six morphologically and ecologically distinct species, the Ursinae (a bear subfamily) species (Kutschera et al., 2014) and the *Geospiza* clade (a genus of Darwin’s finches) (Farrington, Lawson, Clark, & Petren, 2014). We estimated in particular 1) the relative amount of gene flow that has shaped each data-set, and 2) the corresponding average number of ancestral species.

## Material and methods

### Inference method for the GBD-models

When considering several sampled genomes all containing *n* genes, a set of *n* gene trees is obtained for each particular parameter setting and each realization of the model. To characterize a set of gene trees, we employed a multidimensional summary statistic defined as the distribution of pairwise distances between gene trees. Because the GBD-models are time oriented, a tree metric for rooted trees was necessary. Among this class of metrics, we evaluated three metrics: one accounting only for topology, the Robin-son–Foulds (RF) metric (Robinson & Foulds, 1981), and two metrics accounting for both branch lengths and topological differences, the Billera-Holmes-Vogtmann (BHV) metric (Billera, Holmes, & Vogtmann, 2001) and the Kendall and Colijn (KC) metric (Kendall & Colijn, 2016).

The RF metric between two trees, *i.e.* the distance between two trees, is computed as the number of bipartitions of the taxon set congruent with one tree but not with the other tree. The BHV metric is based on a view of tree space as a quadrant complex with quadrants sharing faces. Two trees with the same topology lie in the same quadrant, otherwise they lie in two distinct quadrants. At a common edge between two quadrants, the incongruent internal branches between trees have lengths equal to zero. Then a distance can be calculated between two rooted trees as the shortest path across these interconnected quadrants. The KC metric corresponds to the Euclidean distance between two vectors (two by tree). One vector records the number of edges between the most recent common ancestor and each pair of tips and the second records the path length on the considered edges in the first vector (lengths of each tip are also recorded in the second vector). BHV and KC distances do not rely only on the topology but also on branch lengths. The difference in topology is weighted by the branch lengths supporting these topologies, therefore uncertainties causing polytomies (or a branching pattern close to a polytomy) in gene trees will only marginally affect our results.

To compare trees that did not evolve on the same time scale, RF, BHV and KC distances were computed on re-scaled trees. For each set of gene trees issued from a single simulation or data-set, we rescaled all the trees so that the median of the most recent node depth is 1. After this step, the relative difference in branch lengths remains the same among each set of gene trees.

To estimate model parameters on empirical or test (simulated) trees, we employed the Kullback-Leibler (KL) divergence (package ’FNN’ in R) as a distance metric by minimizing this distance between the distributions of pairwise distances (BHV, KC or RF) of empirical, and test trees, with simulated trees. The lower the KL divergence the better is the fit.

### Inference method accuracy

To define which distance metric was the more suitable, we tested the accuracy of our inference method, detailed above, using either BHV, KC (with the parameter defining the balance between branch lengths and topology set to 0.5) and RF distances.

Using the GBD-backward model, we built gene trees for 204 parameter combinations (with *N* = 6 and *n* = 10) by varying two parameters, *a* and *b*, and fixing *d* = 1 and *c* = 200. The number of time units *t* was set to 5,000. We performed 85 replicates (75 replicates to build the reference distributions and 10 replicates as a test data-set) under each parameter combination in a grid of 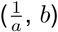 with 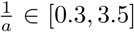, every 0.2, and *b ∈* [0.01, 0.12], every 0.01. For each of the 204 test data-sets, the parameters 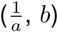 were inferred by minimizing the KL distance between the pairwise distance distribution of the test trees and the pairwise distance distribution (95% confidence interval) of the reference trees from the grid.

Because KC distances showed a better performance in inferring the model parameters (see supplementary results), we used only this metric for the other analyses of the study.

### Comparison of the GBD-models

Next, we aimed to evaluate i) the dynamic of coalescence events between two genomes, called here the coalescence profiles, ii) the number of ancestral species through time for one genome, and iii) the maximal number of gene lineages of one genome located in one (ancestral) species. We performed simulations for *n* = 20 genes, with *α* = 0.1, *β* = 0.2, *δ* = 0.06 and *K* = 30 for the GBD-forward, and *a* = 2, *b* = 0.2, *d* = 6 and *N* = 30 for the GBD-backward model. Additionally, to visually compare the reconstructed genealogies obtained with the GBD-forward and the GBD-backward model we performed simulations for genomes containing *n* = 5 genes, with *α* = 0.5, *β* = 1, *δ* = 0.2 and *K* = 30 for the GBD-forward, and *a* = 1, *b* = 0.1, *d* = 2 and *N* = 10 for GBD-backward model.

Both models gave qualitatively similar results (see Results section). However because the GBD-forward model is computationally prohibitive, all the following analyses were performed with the GBD-backward model. A simulation, with *N* = 6 (*K* = 30 for the GBD-forward to be able to reconstruct genealogies of *N* = 6 genomes), *n* = 10, *a* = *α* = 1, *b* = *β* = 1, *c* = 200 and *d* = *δ* = 1, took about 10 hours for the GBD-forward model and 10 minutes for GBD-backward model (Intel(R) Core(TM) i7-6700 CPU).

### A single sampled genome (GBD-backward model)

To evaluate the variation in the number of ancestral species with the intensity of gene flow, we performed simulations for a single sampled genome containing *n* genes (with *n* = 20, 50, 100, 200), and varied the relative amount of gene flow (erosion rate *d*) compared to genetic differentiation (*non-homologous attraction* rate *a*), ratio 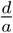 (with *a* = 1 and *d ∈* [0.2, 2], every 0.2). The simulation was run for 10,000 steps. We sampled the number of ancestral species every 500 steps starting at time *t* = 5, 000, and averaged these 11 values for each simulation. For each set of parameters, 5 replicates were performed and averaged.

A model is said to be *sampling consistent* if the same outcome is expected for any *k* sampled genes independently of the total number *n* of genes in the genome. To evaluate the validity of this property, we randomly sampled *k* = 20 genes from each genome of *n ≥* 20 genes and computed their average number of ancestral species.

### A sample of several genomes (GBD-backward model)

We evaluated the influence of the number *n* of genes (with *n* = 10, 15, 20), of the number of species *N* (with *N* = 6, 10), and of the relative amount of gene flow 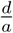 (with *d* = 1 and 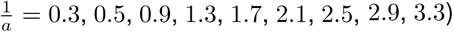, on gene tree diversity (KC distances) (Fig. 6A). The other parameters were fixed, with *b* = 0.05 and *c* = 200.

For the same values of *d* and *c* but with 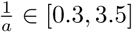, every 0.2, and for *n* = 10, *N* = 6, we also evaluated the influence of the *homologous attraction* rate *b* (with *b* = 0.01, 0.02, 0.05, 0.12) on gene tree diversity (KC distances) (Fig. 6B).

### Inference from empirical data-sets

#### Empirical data-sets

To evaluate if the GBD-backward model correctly reproduces the signal left by gene flow in gene trees we compared empirical gene trees (shaped under gene flow) with simulated gene trees under the GBD-backward model and under the MSC model. The adequacy between the simulated trees and empirical gene trees was estimated by comparing the distributions of pairwise gene tree distances of simulated vs empirical data-sets. The empirical clades have been chosen for their moderate phylogenetic depth, good sampling coverage and known conflicting gene trees. The bear data-set comprised 14 autosomal introns for 6 bear species (*Helarctos malayanus*, *Melursus ursinus*, *Ursus americanus*, *U. arctos*, *U. maritimus*, and *U. thibetanus*) and 2 outgroups (*Ailuropoda melanoleuca* and *Tremarctos ornatus*) (Kutschera et al., 2014). The sequences were downloaded from GenBank (supplementary Table S1). As done by Kutschera et al. (2014), all variation within and among individuals was collapsed into one single 50% majority-rule-consensus sequence for each of the 8 species. The phylogenetic trees were built with the program BEAST v. 1.8.3. (Drummond, Suchard, Xie, & Rambaut, 2012), with the parameters used by Kutschera et al. (2014): Yule prior to model the branching process, strict clock, a normal prior on substitution rates (0.001 *±* 0.001) (mean *±* SD), minimum age of 11.6 My for the divergence of *A. melanoleuca* from other bears (exponential prior: mean = 0.5; offset = 11.6), and 10 million generations with sampling every 1000 generations. The models of DNA evolution were estimated by modeltest (function ‘modelTest’, package ’phangorn’ in R (Schliep, 2011)) (supplementary Table S2). The monophyly of the ingroup and the topology among the outgroups were constrained according to the topology depicted by Kutschera et al. (2014).

The second data-set comprised 7 nuclear markers for 6 finch species (*Geospiza conirostris*, *G. fortis*, *G. fulginosa*, *G. magnirostris*, *G.scandens*, and *G. septentrionalis*) and 2 outgroups (*Camarhynchus psittacula* and *Platyspiza crassirostris*) (Farrington et al., 2014). The sequences were downloaded from GenBank (supplementary Table S3). The phylogenetic trees were built with the program BEAST v. 1.8.3. (Drummond et al., 2012) with the parameters used by (Farrington et al., 2014): coalescent constant size prior to model the branching process, strict clock, substitution rate equal to 1, specific models of DNA evolution defined by the authors (supplementary Table S2), and 10 million generations with sampling every 1000 generations. The monophyly of the ingroup and the topology among the outgroups were constrained according to the topology depicted by Farrington et al. (2014).

#### Estimation of parameters under the multi-species coalescent (MSC) model

We optimized the MSC model for *N* = 6 species by varying two parameters, the speciation rate *λ* and the extinction rate *µ*, and fixing the coalescence rate to 1. Birth-death trees of 6 tips (function ’sim.bdtree’, package ’geiger’ in R) were simulated in a grid of (*λ*, *µ* = *mλ*) with *λ ∈* [0.02, 0.34], every 0.02, and *m ∈* [0.1, 0.65], every 0.05. Because we simulated small trees (6 tips), the degree of variation between trees simulated with the same parameters was high. Therefore for each value of (*λ*, *µ*) we randomly selected 75 species trees for which the crown age did not differ by more than 2.5% from the expected crown age. Next, we simulated 10 gene genealogies for each species tree (coalescence rate fixed to 1).

If the diversification rate (speciation rate minus extinction rate) is low, all the homologous genes will coalesce before the next node in the species tree, so that all the gene trees will have the same topology. On the contrary, if the diversification rate is too fast, some homologous genes will not have time to coalesce before the next node of the species tree, resulting in incongruent gene trees due to the randomness of coalescences (ILS).

#### Estimation of parameters under the gene-based diversification (GBD-backward) model

Equivalently, we optimized the GBD-backward model for *N* = 6 by varying two parameters, here *a* and *b*, and fixing *d* = 1 and *c* = 200 (recall *c* is given a sufficiently large value that coalescences are instantaneous). Since increasing *n* has no effect on KC distances (see results and Fig. 6), we simulated genomes with *n* = 10 genes. Each simulation was conducted until the coalescence of all homologous genes. We performed 75 replicates under each parameter combination in a grid of 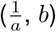 with 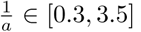, every 0.2, and *b ∈* [0.01, 0.12], every 0.01.

For both models (MSC and GBD-backward) we employed the Kullback-Leibler (KL) divergence (package ’FNN’ in R) as a distance metric to find the best set of parameters by minimizing this distance between the distributions of KC pairwise distances of empirical and simulated trees. The lower the KL divergence is the better is the fit.

## Results

### Inference method accuracy

Using simulated data-sets we showed that our inference method, comparison of the distribution of KC pairwise distances among a set of trees, was able to give reliable estimates of simulated parameters despite its simplicity (see supplementary results figure S1). This simple inference method is sufficient to estimate the parameters of the model having supposedly shaped the gene trees of the data set. More subtle methods will be developed in the future to account for more complex features, such as differential gene flow depending on putative gene categories, and to infer the very history of the embedding of gene lineages into species.

### Comparison of the GBD-models

Even if the two models, GBD-forward and GBD-backward, are not rigorously the inverted image of one another, they showed a qualitatively similar pattern in gene genealogies and in the distribution of gene lineages among species through time (Figs. 3 and 4).

**Figure 3:**
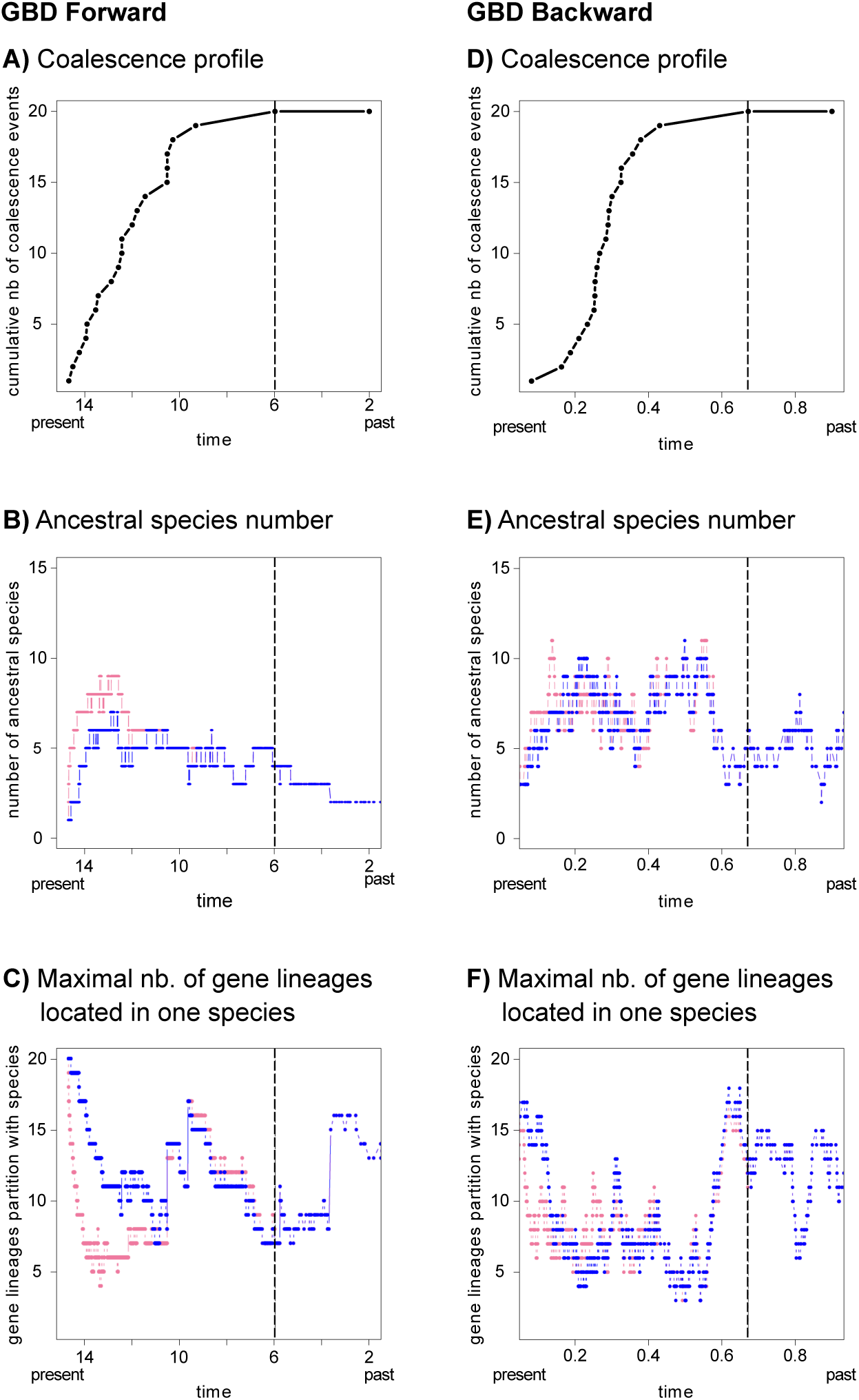
Comparison of the GBD models. A) and D) Coalescence profiles of two genomes sampled at present time. B) and E) The corresponding number of ancestral species through time for each genome. C) and F) The maximal number of gene lineages belonging to each genome located in one species. The dashed line represents the last coalescence events between the two sampled genomes. Simulations were performed for *n* = 20 genes, with *α* = 0.1, *β* = 0.2, *δ* = 0.06 and *K* = 30 for the GBD-forward, and *a* = 2, *b* = 0.2, *d* = 6 and *N* = 30 for the GBD-backward model. Nb.: number.

**Figure 4:**
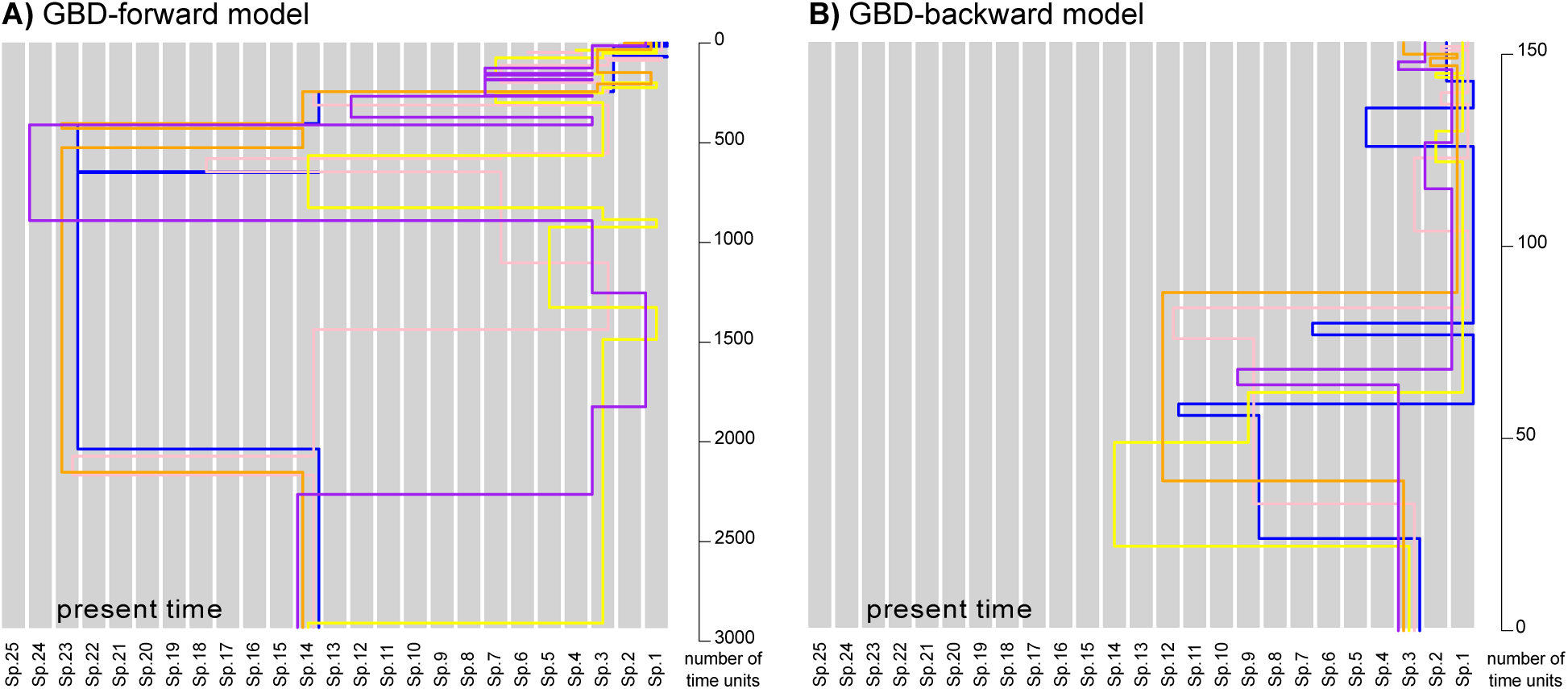
Genealogies of a single genome generated with the GBD-forward (A) and GBD-backward models (B). The labels/locations of species (or populations, depending on the point of view, retrospective *vs* prospective) are neutral. A) Parameter settings: *α* = 0.5, *β* = 1, *δ* = 0.2, *n* = 5 and *N* = 30. B) Parameter settings: *a* = 1, *b* = 0.1, *d* = 2, *n* = 5 and *N* = 10.

The coalescence profiles (*i.e.* the cumulative number of coalescence events through time) were similar in shape for both models (Figs. 3 and S2) with a rapid increase followed by a plateau until the last coalescence event. However, contrary to the GBD-backward model, the GBD-forward model can generate simultaneous coalescences (as in our simulation, Fig. 3A). If gene flow is too weak relatively to the other parameters, some of the simultaneous coalescences modelled at a colonization event will remain together (*i.e.* happening at the same time *t*) and appear as a multifurcation in reconstructed trees.

The number of ancestral species through time was also comparable between the two models (Fig. 3B and E). Going backward in time, after a first increase in the number of ancestral species, gene lineages tend to be brought back together in fewer ancestral species whether it is due to non-homologous attraction in the GBD-backward model or because genes were co-adapted at this time in the GBD-forward model. In the GBD-backward model, once at least a pair of homologous genes have coalesced, the two genomes will be attracted toward this coalesced gene lineage increasing the probability for two other homologous genes to be in the same species and thus to coalesce. The attraction will be even stronger when many genes have coalesced, explaining the latter decrease in the number of ancestral species (Fig. 3E).

In the GBD-forward model, going forward in time, the number of ancestral species increases and suddenly decreases. Just after a *colonization* event, the new species can easily exchange genes with related species. Gene lineages will then acquire new mutations, isolating each genome from the others, explaining the rapid decrease in the number of ancestral species toward the present (Fig. 3B). Note that in both cases, going backward in time, once all the homologous genes have coalesced they follow the exact same path through species (the blue and pink lines are confounded in the Fig. 3B-F).

Moreover, because they are co-adapted, genes sampled in the same species at present time should have spent time together in the same species more often than by chance in the past. This property was indeed observed in both models, with genes sampled at present time frequently found together in the same species in the past (see Figs. 3C, F and 4), where at least five gene lineages (over 20) lie in the same ancestral species at all times).

### A single sampled genome (GBD-backward)

With *N* = 1 sampled genome containing *n* genes, we let *A*(*t*) = (*A*_1_(*t*)*, …, A_n_*(*t*)) denote the sorting of genes into ancestral species *t* units of time before the present. More precisely, *A_k_*(*t*) denotes the number of ancestral species containing *k* gene lineages, so that 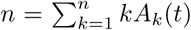 and 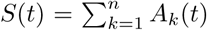 is the total number of species at *t* ancestral to the sampled genome. For each *ε ∈* (0, 1], we will also be interested in the number 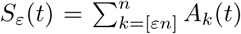 of ancestral species containing at least a fraction *ε* of the genome (with [*x*] denoting the smallest integer larger than *x*). All stationary quantities will be denoted by the same symbols, replacing *t* with *∞*.

We will call a *block* at (backward) time *t* a (maximal) set of gene lineages that lie in the same species at time *t*. The transition rates can be specified as follows in terms of the configuration of gene lineages into blocks (*i.e.*, ancestral species). For each pair of blocks containing (*j, k*) lineages, non-homologous attraction occurs at rate *ajk* and results in the configuration (*j −* 1*, k* + 1). For each block containing *j* lineages, erosion occurs at rate *dj* and results in the block losing one lineage; simultaneously a new block containing 1 single lineage is created. These are exactly the same rates as in the well-known Moran model with mutation under the infinite-allele model (Moran, 1958), replacing ‘block’ with ‘allele’, ‘attraction’ by ‘resampling’ (simultaneous birth from one of the *j* carriers of a given allele and death of one of the *k* carriers of another given allele) and ‘erosion’ with ‘mutation’ (mutation appearing in one of the *j* carriers of a given allele into a new allele never existing before). For this Moran model,

- the total population size is *n*;
- at rate *a* for each oriented pair of individuals independently, the first individual of the pair gives birth to a copy of herself and the second individual of the pair is simultaneously killed;
- mutation occurs at rate *d* independently in each individual lineage.

As a consequence, *A*(*t*) has the same distribution as the allele frequency spectrum in the Moran model with total population size *n*, resampling rate *a* and mutation rate *d*, starting at time *t* = 0 from a population of clonal individuals (one single block). In particular, the distribution of *A*(*∞*) is the stationary distribution of the allele frequency spectrum, which is known to be given by Ewens’ sampling formula with scaled mutation rate *d/a* (Durrett, 2008; Ewens, 1972; Ewens & Tavaré, 2006). Expectations of this distribution are:

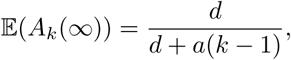

so that

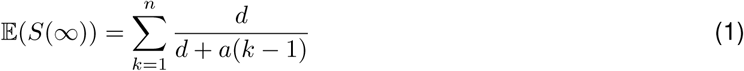

and

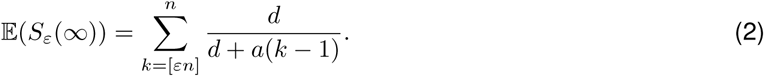

In particular, as *n → ∞*,

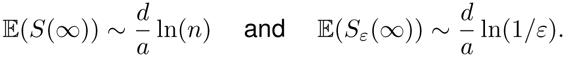

At stationarity, and particularly for large values of 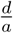, the mean number of ancestral species *S*(*∞*) obtained from simulations was equal to the mathematical prediction (figure 5A). In particular, the mean number of ancestral species at stationarity increases with 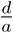.

**Figure 5:**
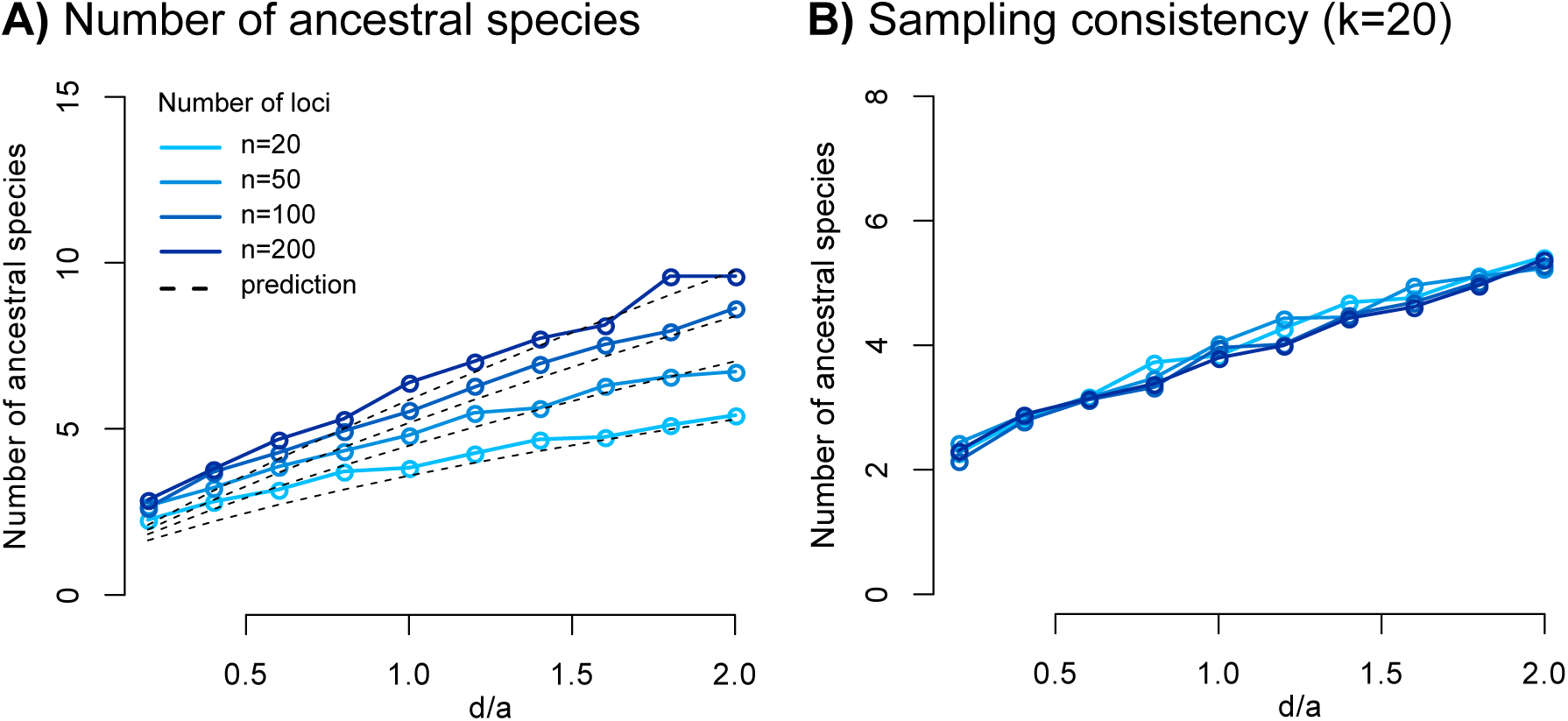
Evaluation of the GBD-backward model for a single sampled genome with *n* genes. Parameter settings: *a* = 1, *d ∈* [0.2, 2], every 0.2, and *n* = 20, 50, 100, and 200. The number of time units *t* was set to 10, 000. We sampled the number of ancestral species every 500 time units starting at time *t* = 5, 000, and averaged them for each simulation. For each set of parameters, 5 replicates were performed and averaged. A) Number of ancestral species depending on the number of genes *n* and on the ratio 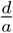, for one sampled genome. B) To assess the sampling consistency of our models, *k* lineages were randomly sampled. The number of ancestral species reported is the number of ancestral species of these *k* genes only.

An additional key feature of this model is *sampling consistency*. In words, the history of a sample of *k* genes taken from a genome of *n* genes does not depend on *n*. This property can again be deduced from the representation of our model in terms of the better known Moran model. Indeed, the dynamics of a sample of *k* individuals in the Moran model does not depend on the population size, as can be seen from the so-called lookdown construction (due to P. Donnelly and T. Kurtz and clearly exposed by Etheridge 2011). The simulations performed with *k* genes randomly sampled from each genome of *n* genes, are in agreement with this claim of sampling consistency: the number of ancestral species at stationarity 𝔼(*S_ε_*(*∞*)) is independent of the number of genes *n* (Fig. 5B).

### A sample of several genomes (GBD-backward)

Using simulations, we evaluated the GBD-backward model for several sampled genomes (*N >* 1) under several combinations of parameters. As expected, gene tree variation, measured by KC distances, increased with 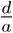, *i.e.* the relative amount of gene flow, and with the number of species *N* . Conversely our results showed that the number of genes *n* had no effect on distances (Fig. 6A). This last result, the lack of influence of *n* on gene tree variation, is of particular interest, because one usually has only access to a fraction of a genome. It shows that regardless of the number of genes sampled, the resulting gene tree variation will remain the same as long as gene trees have been shaped by processes with similar parameter values.

**Figure 6:**
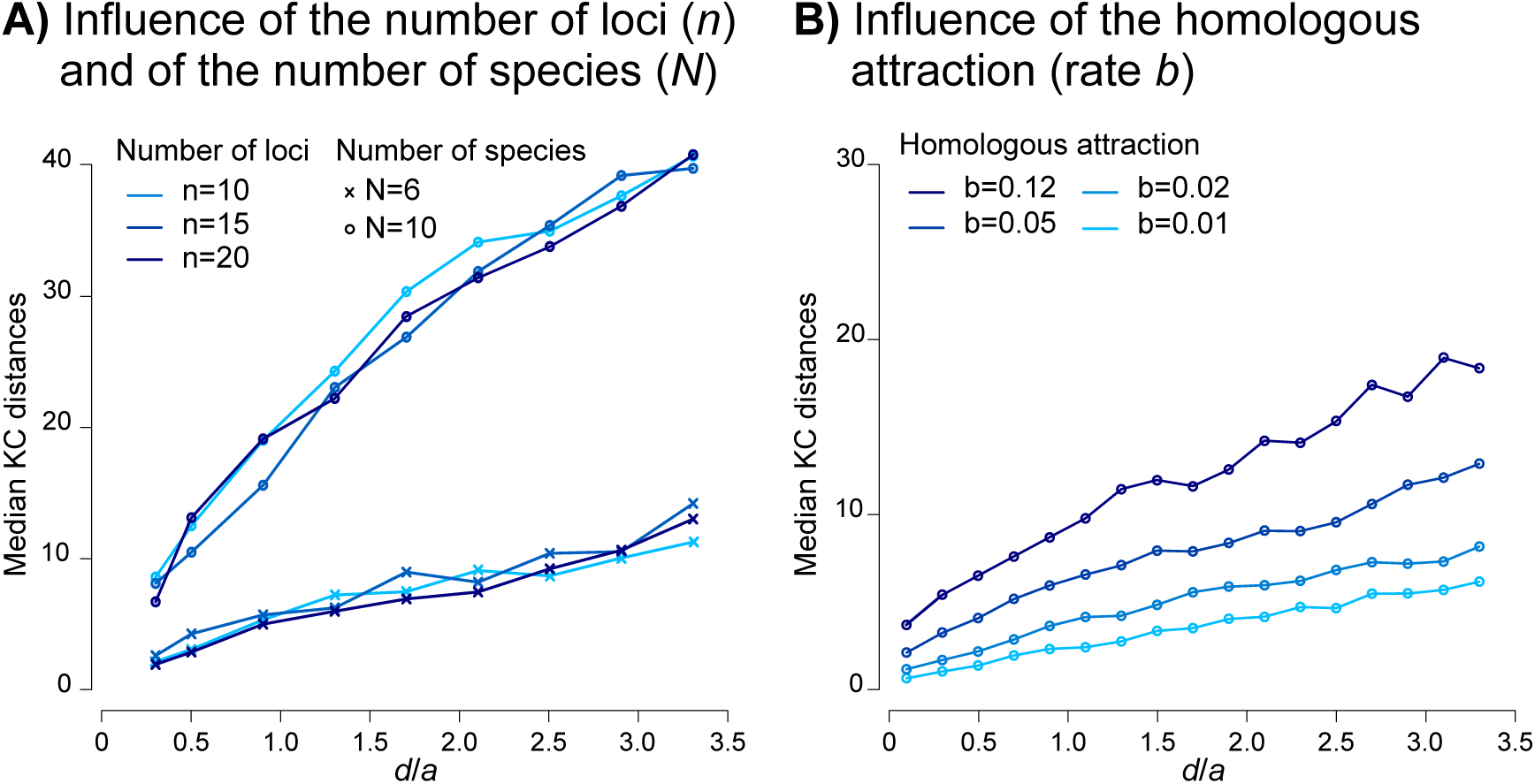
Kendall-Colijn (KC) distances among sets of gene trees simulated under the gene-based diversification (GBD-backward) model. For each set of parameters, with *t* = 5, 000 (enough to reach the coalescence of all homologous genes), the median KC distances were calculated. A) Influence of the number of genes *n* (with *n* = 10, 15, and 20), of the number of species *N* (with *N* = 6 and 10), and of the ratio 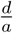 on the KC distances. Parameter settings: *b* = 0.05, *d* = 1, *c* = 200, and 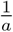 = 0.3, 0.5, 0.9, 1.3, 1.7, 2.1, 2.5, 2.9, 3.3. For each set of parameters, 7 simulations were performed. B) Influence of the *homologous attraction* rate *b* and of the erosion-to-non-homologous attraction ratio 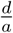 on the KC distances. Parameter settings: *n* = 10, *N* = 6, *b* = 0.01, 0.02, 0.05, 0.12, *d* = 1, *c* = 200, and 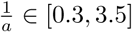, every 0.2. For each set of parameters, 75 simulations were performed.

Our results also showed that as the *homologous attraction* rate *b* decreases, and for the same value of 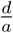, gene trees were more similar (lower KC distances) (Fig. 6B). When a long period of time elapses between two homologous attraction events (low *b*), all the genes belonging to the two genomes that have started to coalesce, have enough time to be attracted toward the same ancestral species, and thus coalesce before the next homologous attraction event, in spite of gene flow.

### The GBD-backward model correctly captures the signal left by gene flow in empirical data-sets

To estimate model parameters, we minimized the Kullback-Leibler (KL) divergence between the distributions of KC pairwise distances of empirical and simulated trees (Fig. 7). Under the multi-species coalescent (MSC) model, the most likely parameter values were *µ* = 0.2 *× λ* and *λ* = 0.08 (KL divergence = 0.39) for the bears and *µ* = 0.60 *× λ* and *λ* = 0.20 (KL divergence = 0.19) for the finches. Under the gene-based diversification (GBD-backward) model, the most likely parameter values were *b* = 0.03 and 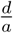 = 2.1 (KL divergence = 0.24) for the bears and *b* = 0.08 and 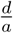 = 1.7 (KL divergence = 0.01) for the finches (Fig. 7). We noted longer tailed distributions for the distances between trees modeled under the MSC model than for the empirical data-sets (Fig. 8). This skewed distribution obtained with the MSC model explains why we did not detect a sharp peak in the optimization landscape for the MSC model (Fig. 7).

**Figure 7:**
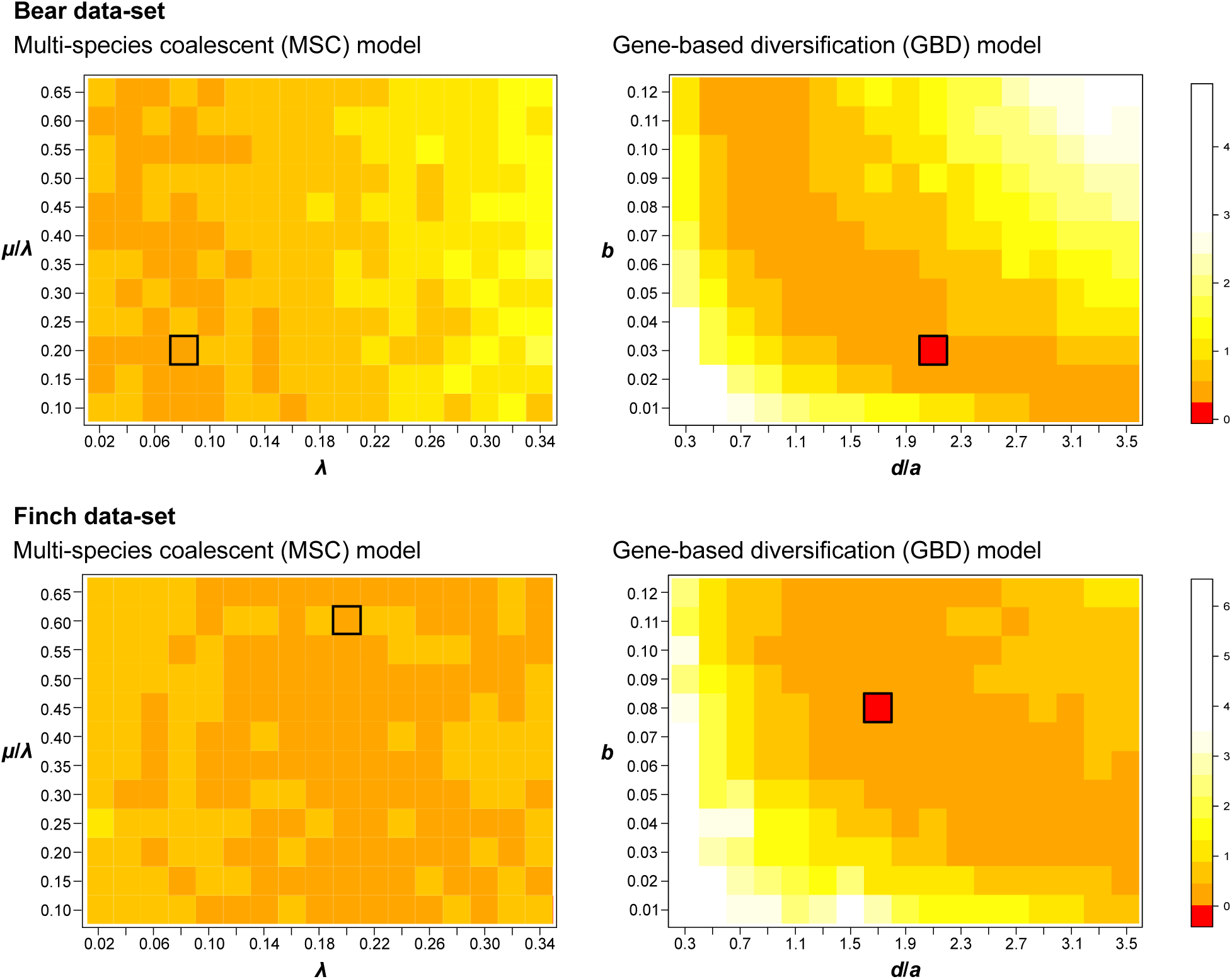
Minimization of the Kullback-Leibler (KL) divergence between empirical and simulated trees, *i.e.* between their distributions of KC pairwise distances. Two parameters were optimized for each model. The *speciation* rate (*λ*) and the *extinction* rate (*µ*) for the multi-species coalescent (MSC) model (with coalescence rate set to 1). The *homologous attraction b* and the ratio of the *erosion* rate over the *non-homologous attraction* rate (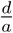) for the gene-based diversification (GBD-backward) model (with *d* set to 1). For each set of variables, 75 simulations were performed and averaged. The same color scale was used for each empirical data-set. For each optimization analysis, the cell for which we found the best fit between empirical and simulated trees (smallest KL divergence) is framed.

**Figure 8:**
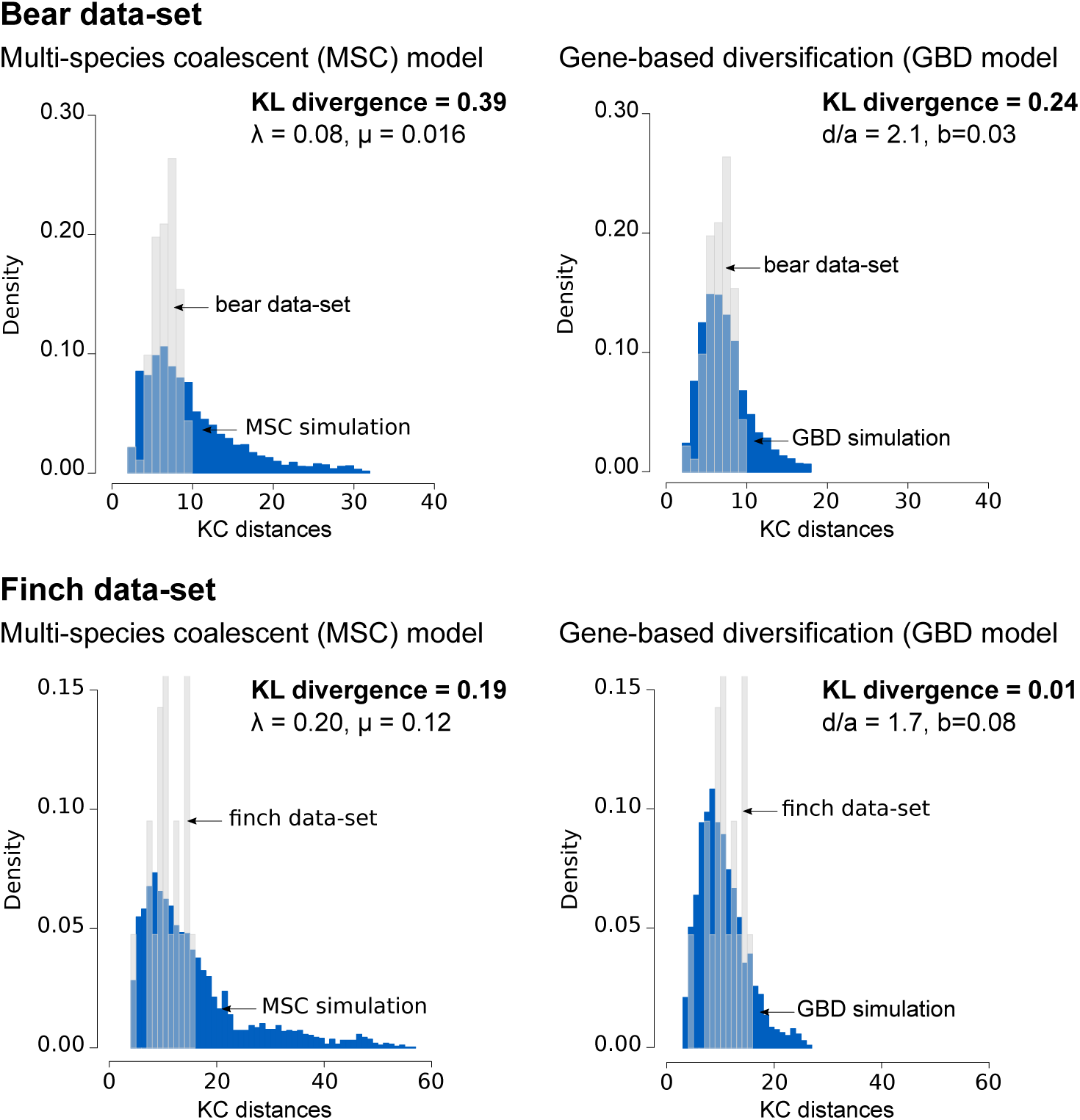
Best fit between empirical and simulated trees, *i.e.* between their distributions of KC pairwise distances (selected cells of Fig. 7). For each set of variables, 75 simulations were performed and averaged. *a*: *non-homologous attraction* rate, *b*: *homologous attraction* rate, *d*: *erosion* rate (set to 1), *λ*: *speciation* rate, *µ*: *extinction* rate, KL: Kullback-Leibler.

However, for both data-sets, the selected simulated distribution (GBD model) and the empirical distribution do not match perfectly. This difference, an excess of intermediate distances and a lack of large distances among empirical trees could reflect the exclusion of too incongruent genes when the data-sets were built, and/or the presence of functionally or physically linked genes (preferentially found together in ancestral species).

Comparing the parameters *λ* and *µ* to *b* and 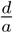 is not straightforward as the two models, MSC and GBD-backward, are built under different assumptions. However in both cases, the parameters influence the diversity among trees (shape of the distribution of KC pairwise distances). A greater variation among trees is expected with increasing *λ* and decreasing *µ*, and with increasing 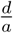 and *b*, allowing us to explore the parameter landscape to find the setting that minimizes the distance between simulations and empirical data-sets for each model.

Given our results and the mathematical predictions, the time-averaged number *S_ε_*(*∞*) of ancestral species to the sampled genome containing at least 10% of the genome (*ε* = 0.1) when *n → ∞* is 4.8 for the bear data-set and 3.9 for the finch data-set.

## Discussion

Within species, gene flow allows the maintenance of species cohesion in the face of genetic differentiation (Morjan & Rieseberg, 2004; Slatkin, 1987), preventing genetic isolation of populations and the subsequent emergence of reproductive barriers leading to speciation (Coyne & Orr, 2004). Among species, the existence of gene flow challenges the notion of a species genealogy as well as the current concepts of species. Indeed, if gene flow is as pervasive as recent empirical studies suggest (Clark & Messer, 2015; Cui et al., 2013; Gallus et al., 2015; Jónsson et al., 2014), the genealogical history of species should be represented as a phylogenetic network encompassing the mosaic of gene genealogies. Similarly, it seems very conservative to delineate species based on the widely used biological species concept (reproductive isolation) (Mayr, 1942), or phylogenetic species concept (reciprocal monophyly) (Papadopoulou et al., 2008). Because of the ubiquity of gene flow, which can persist for several millions of years after the lineages have started to diverge (*i.e.*, onset of speciation) (Bolnick, Near, & Noor, 2005; Mallet, 2005), species should be rather defined by their capacity to coexist without fusion in spite of gene flow (Mallet, 2008; Samadi & Barberousse, 2006).

The simplified view of diversification, consisting in representing lineages splitting instantaneously into divergent lineages with no interaction (gene exchange) after the split, has been preventing evolutionary biologists from fully apprehending diversification at the genomic level and from correctly interpreting discrepancies between gene histories. Indeed, conflicting gene trees make the interpretation of their evolutionary history difficult. However, we argue that phylogenetic incongruence among gene trees should not be considered as a nuisance, but rather as a meaningful biological signal revealing some features of the dynamics of genetic differentiation and of gene flow through time and across clades. Current phylogenetic methods rely on the assumption that gene trees are constrained within the species tree, and that gene flow occurs infrequently between species. For many data-sets such as sequence alignments of genomes sampled from young clades, such methods could lead to an evolutionary misinterpretation of gene trees, and in the worst case to species trees with high node support while the gene trees had very different evolutionary histories (Long & Kubatko, 2018). These observations urge for a change of paradigm, where gene flow is fully part of the diversification model. To consider the ubiquity of gene flow across the Tree of Life and its broad effect on genomes described by many recent studies, we have developed a new framework focusing on gene genealogies and relaxing the constraints inherent to the MSC paradigm. This framework is implemented in a mathematical model that we named the gene-based diversification (GBD-forward) model. We have also developed a complementary version of this model, the GBD-backward model, speeding up the simulations thanks to a coalescent approach.

### The GBD-backward model

Under the GBD-backward model, gene genealogies are governed by four parameters: non-homologous attraction rate *a*, homologous attraction rate *b*, coalescence rate *a*, and erosion rate *d* (Fig. 2).

*Non-homologous attraction* models genetic differentiation resulting in reproductive isolation. The slower genes accumulate mutations and differentiate, the more time can be spent by gene lineages in different species. Hence when genomes differentiate slowly, the rate of non-homologous attraction is low. *Homologous attraction* corresponds to finding the most recent common ancestor of the two species at the genomic level. The time spent between homologous attraction events depends crucially on the (phylogenetic distance of the) species sampled at the present. *Coalescence* is in direct correspondence with genetic drift in the GBD-forward model and *erosion* with gene flow.

Each of these parameters influences differently the resulting tree variation, *i.e.* the distribution of the KC distances among trees, that we used here as a summary statistic. Instead of focusing on the main phylogenetic signal alone as done by the current phylogenetic methods, the GBD-backward model makes use of the whole signal generated by all gene trees.

Higher amount of gene flow (large *d* values) and reduced time to untangle gene genealogies before the connection of two other genomes (large *b* values) increase variation among trees. Conversely, when homologous genes coalesce faster (large *c* values) and genes are recalled faster toward the species harboring the other genes of their genome (large *a* values) gene trees are expected to be more similar.

After evaluating this model under various sets of parameters, we applied it to analyze two empirical multilocus data-sets for which gene tree conflicts obscure the evolutionary history.

### Gene flow among bears and among finches

In many cases, such as among bears and finches, gene flow is frequent and complicates the relationships between species, challenging the notion of a unique species tree. A strictly bifurcating lineage-based model will not adequately reflect those complex evolutionary patterns. On the contrary, models developed under the *genomic view of diversification* framework, *i.e.* relaxing species boundaries and accounting for gene flow, will better reproduce the complex history of gene genealogies under pervasive gene flow. Note that we considered a simple scenario with no ILS and statistically exchangeable genes resulting in a model with only three parameters, but given the simplicity and the flexibility of our model, many extensions may be considered to address scenarios that could not have been considered previously, opening up new perspectives in the study of speciation and macro-evolution.

Our results showed support for the hypothesis that gene flow has shaped the gene trees of bears and finches (Fig. 8). For the bear data-set, we found that each species had on average in the past about 4.8 ancestral species carrying at least 10% of its present genome (Equation (2)). This result is in line with previous studies reporting gene flow between pairs of bear species (Cahill et al., 2013; Hailer et al., 2012; Kutschera et al., 2014; S. Liu et al., 2014; Miller et al., 2012). Moreover, a recent phylogenomic study (869 Mb divided into 18,621 genome fragments) confirmed the existence of gene flow between sister species as well as between more phylogenetically distant species (Kumar et al., 2017). The authors used the *D*-statistic (gene flow between sister species) and *D_FOIL_*-statistic (gene flow among ancestral lineages, Pease and Hahn 2015) to detect gene flow among the 6 bear species. Using their results, for each pair of species *ij* among the *N* species, we determined if the species *j* has contributed (*g_ij_* = 1) or not (*g_ij_* = 0) to the genome of the species *i* (with *g_ii_* = 1), and calculated the average number of ancestral species 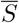 as follows:

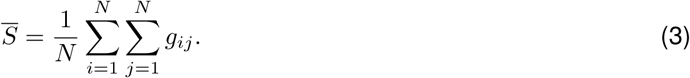

Using this equation on the results of the phylogenomic study (Kumar et al., 2017), we found on average 5.3 ancestral species for each of the Ursinae bears, close to the estimate of 4.8 obtained with the GBD-backward model.

We detected lower gene flow among finches than among bears. Each finch species had on average in the past 3.9 ancestral species (for the subsample of gene trees analyzed here), which is also consistent with the extensive evidence that many species hybridize on several islands (Freeland & Boag, 1999; P. R. Grant & Grant, 1997; P. R. Grant, Grant, & Petren, 2005; Sato et al., 1999). Because of gene flow very little genetic structure was detected by a Bayesian population structure analysis, (only 3 genetic populations among the 6 *Geospiza* species, see Farrington et al. 2014). Each of the 2 species *G. magnirostris* and *G.scandens* was mostly characterized by a single genetic population, therefore had about 1 ancestral species each. Conversely 4 *Geospiza* species shared the same genetic population, suggesting 4 ancestral species for each of these 4 species. Taking together these results roughly indicate that each of the 6 *Geospiza* species had in average 3 ancestral species, which is slightly lower than the GBD-backward estimate of 3.9.

### Perspectives

Phylogenetic models and methods inferring macro-evolutionary history, such as speciation and extinction rates, trait evolution or ancestral characters, have become increasingly complex (Morlon, 2014; Pyron & Burbrink, 2013; Stadler, 2013b). Yet, the raw material used by these methods is often reduced to the species tree, which can be viewed as a summary statistic of the information contained in the genome. We argue here that a valuable amount of additional signal, not accessible in species trees, is contained in gene trees, and is directly informative about the diversification process. Indeed, because genetic differentiation and gene flow impact each gene differently, genes may have experienced very different evolutionary trajectories. In order to make use of the entire information conveyed by gene trees, we have proposed here a new approach to study diversification, the genomic view of diversification, under which gene trees shape the species tree rather than the opposite. This approach aims at better depicting the intricate evolutionary history of species and genomes. We hope that this view of diversification will contribute to pave the way for future developments in the perspective of inferring diversification processes directly from genomes rather than from their summary into one single species tree. One of the challenges in this direction will be to propose finer inference methods than the simple, but reasonably satisfactory, method used here, based on a single multidimensional summary statistic, the distribution of pairwise KC distances between gene trees.

## Supporting information

Supplementary results

GBD code

## Supplementary material

The code for the models is available as Supplementary Material.

## Acknowledgments

The authors thank the *Center for Interdisciplinary Research in Biology* (Collège de France, CNRS) for funding. JM is funded by LabEx MemoLife, project *Genomics of Diversification*. The authors also thank the INRA MIGALE bioinformatics platform (http://migale.jouy.inra.fr) for providing computational resources.

