## Supplementary results for "The genomic view of diversification"

We evaluated our inference method, comparison of the distribution of pairwise distances among a set of trees, for three distance metrics (figure S1). Two metrics take into account both branch lengths and topology (Billera-Holmes-Vogtmann (BHV) ans Kendall Colijn (KC)) and one metric (Robinson–Foulds (RF)) only topological differences. The distance metric associated with the better inference results was determined by computing mean squared error (MSE) scores, defined as the average squared difference between the observed (inferred parameters) and predicted values (simulated parameters). Inferences using RF distances performed very poorly, with an average MSE of $2.13$ for the parameter $\frac{d}{a}$ and $2.21\mathrm{e}{-03}$ for the parameter $b$. Better results were retrieved for BHV distances, with an average MSE of $0.68$ for the parameter $\frac{d}{a}$ and $1.39\mathrm{e}{-03}$ for the parameter $b$. KC distances showed a similar performance than BHV distances for the parameter $b$ with an average MSE of $1.58\mathrm{e}{-03}$, but a better performance for the parameter $\frac{d}{a}$ with an average MSE of $0.61$.

Despite low MSE for the parameter $b$ due to simulated values $< 1$, the plots showed inconsistencies for the inference on $b$. This parameter, *homologous attraction*, has only a small influence on gene genealogies as it solely governs the first coalescence event between two genomes. Following this event, the other parameters $a$ and $d$ (*non-homologous attraction* and *erosion* respectively) will govern gene genealogies, and therefore will be more accurately estimated. It should also be noted that the parameter $b$ is not used to estimate the number of ancestral species.

Figure S1: Inference performance on $\frac{d}{a}$ (A) and $b$ (B) using either Billera-Holmes-Vogtmann (BHV), Kendall Colijn (KC) or Robinson–Foulds (RF) distances. Inferences were performed for each of the $204$ parameter combination in a grid of ($\frac{1}{a}$, $b$) with $\frac{1}{a} \in [0.3, 3.5]$, every $0.2$, and $b \in [0.01, 0.12]$, every $0.01$ (with for each parameter combination $75$ replicates to build the reference distributions and $10$ replicates as a test data-set). The parameters ($\frac{1}{a}$, $b$) were inferred by minimizing the KL distance between the pairwise distance distribution of the test trees and the pairwise distance distribution ($95\%$ confidence interval) of the reference trees. The plots encompass $95\%$ of the inferred parameters distribution. Red dots represent the simulated values and black dots the medians of the inferred parameters distributions.

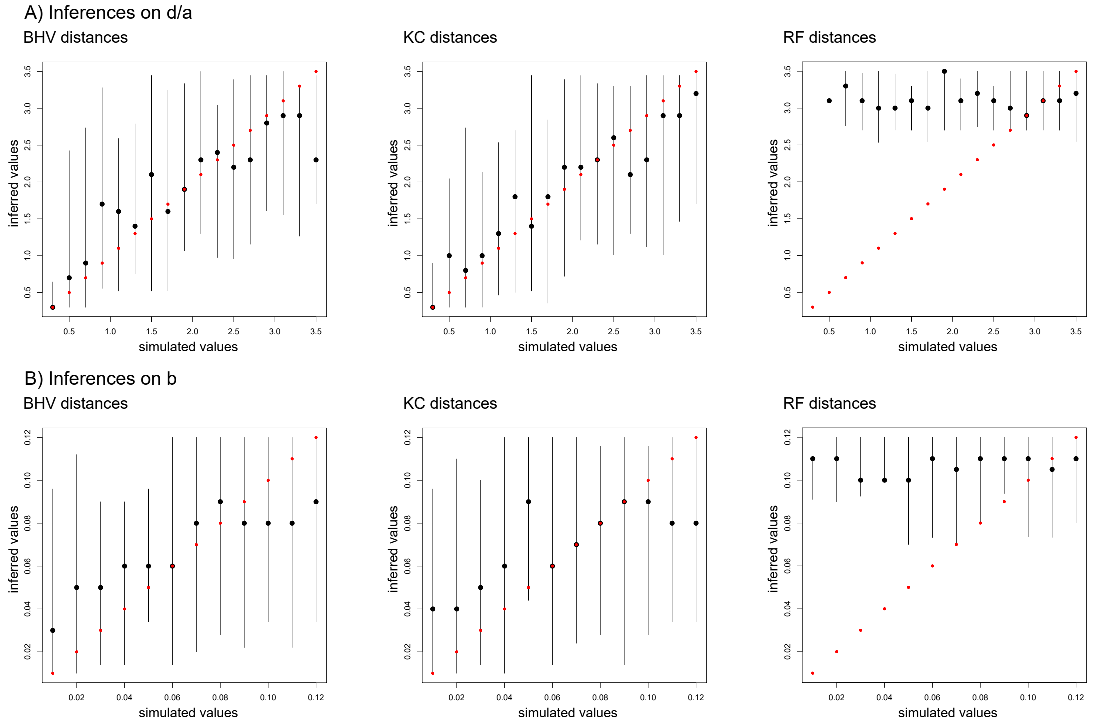

Figure S2: Coalescence profiles of $20$ replicates of the GBD-forward (A) and backward (B) model are plotted in light gray. The black line represent the mean coalescence profile over the $20$ replicates.

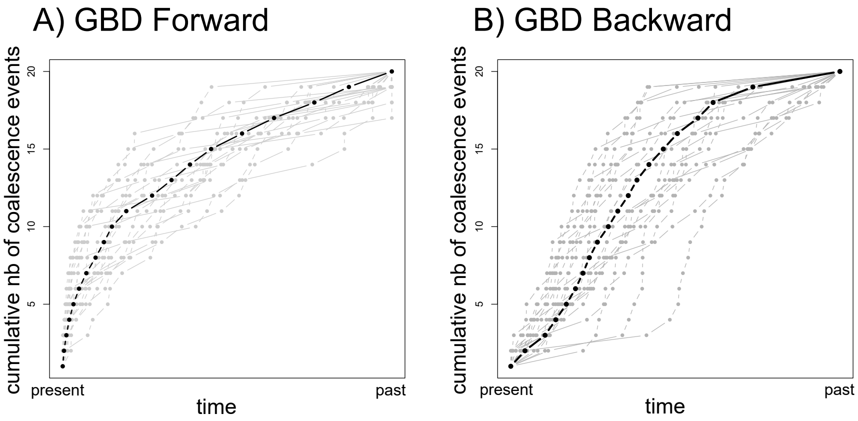
