## Supplementary material for "The genomic view of diversification": GBD code: gbd.backward.R

```
gbd.b <- function(SpNb, LociNb, a, b, c, d, t){

Sp <- SpNb
N <- LociNb
C <- a
C_inter <- b
Tc <- c
D <- d
Tg <- t

tabs <- matrix(integer(Sp*N), nrow = Sp, ncol = Sp*N)
row.names(tabs) <- paste("tab",1:Sp,sep="")
for (i in 1:Sp){
	tabs[i,i] <- N}

locusiN <- matrix(c(sort(rep(1:N,length(tabs[1,]))),c(1:N)), nrow = Sp, ncol = (dim(tabs)[2]*N+N),byrow = TRUE)
row.names(locusiN) <- paste("locusi",1:Sp,sep="")

emptySp <- which(tabs[1,]==0)
for (i in 2:dim(tabs)[1]) {
		emptySp <- intersect(emptySp,which(tabs[i,]==0))}

coal <- matrix(rep(c(1:N),dim(combn(Sp, 2))[2]),nrow = dim(combn(Sp, 2))[2],byrow=TRUE)
row.names(coal) <- paste(combn(Sp,2)[1,],combn(Sp,2)[2,],sep="")

tabCoal <- integer(0)
ssC <- integer(0)
times <- integer(0)
coalT <- integer(0)
coalPair <- integer(0)
CompatL <- 1:N

for(i in 1:Tg) {

## Locations
corrCi <- integer(0)
for (k in 1:Sp){
	corrCx <- integer(0)
	for (i in 1:length(which(tabs[k,]>0))){
		rg <- which(tabs[k,]>0)
		corrCx <- c(corrCx,rep(rg[i],tabs[k,rg[i]]*length(tabs[k,])))}
corrCi <- rbind(corrCi,corrCx)}

corrDi <- integer(0)
for (k in 1:Sp){
	corrDx <- integer(0)
	for (i in 1:length(which(tabs[k,]>0))){
		rg <- which(tabs[k,]>0)
		corrDx <- c(corrDx,rep(rg[i],tabs[k,rg[i]]))}
corrDi <- rbind(corrDi,corrDx)}

corrCDi <- cbind(corrCi,corrDi)

corrCj <- matrix(rep((1:length(tabs[1,])),N), nrow = Sp, ncol = (dim(tabs)[2]*N),byrow = TRUE)
corrDj <- matrix(rep(min(emptySp),N), nrow = Sp, ncol = N,byrow = TRUE)
corrCDj <- cbind(corrCj,corrDj)

## Non-homologous attraction (rate a)

probCi <- integer(0)
for (k in 1:Sp){
	probCx <- integer(0)
	for (i in 1:length(which(tabs[k,]>0))) {
		rg <- which(tabs[k,]>0)
		tab1i <- tabs[k,]
		tab1i[rg[i]] <- 0
		probCx <- c(probCx,rep(C*(tab1i),tabs[k,rg[i]]))}
probCi <- rbind(probCi,probCx)}

## Erosion (rate d)
probDi <- integer(0)
for (k in 1:Sp){
	probDx <- integer(0)
	for (i in 1:length(which(tabs[k,]>0))) {
		rg <- which(tabs[k,]>0)
		probDx <- c(probDx,rep(D,tabs[k,rg[i]]))}
	probDx[which(match(corrDi[k,],(which(tabs[k,]==1)))>0)] <- 0   
probDi <- rbind(probDi,probDx)}

probCD <- cbind(probCi,probDi)

## Homologous attraction (rate b)
for (m in 1:Sp){
	pos1 <- integer(0)
	for (i in 1:length(CompatL)){
		for (k in (1:Sp)[-m]){
			destination1_2 <- unique(corrCDi[k,which(locusiN[k,]== CompatL[i])])
			pos1 <- c(pos1,intersect(which(corrCDj[k,]==destination1_2),which(locusiN[m,]==CompatL[i])))}}
	probCD[m,pos1] <- probCD[m,pos1]+C_inter }

## Coalesced loci
tabsC <- tabs
for (i in 1:dim(coal)[1]){
	i1 <- combn(Sp,2)[1,i]
	i2 <- combn(Sp,2)[2,i]
	cc1 <- which(is.na(coal[i,]))
		if (length(cc1)>0){
		for (k in 1:length(cc1)){
			if (tabsC[i2,unique(corrCDi[i2,][which(locusiN [i2,]== cc1 [k])])]>0) {
			tabsC[i2,unique(corrCDi[i2,][which(locusiN [i2,]== cc1 [k])])] <- tabsC[i2,unique(corrCDi[i2,][which(locusiN [i2,]== cc1 [k])])]-1}}}}

## Remove values for coalesced loci
for (i in 1:dim(coal)[1]){
	i1 <- combn(Sp,2)[1,i]
	i2 <- combn(Sp,2)[2,i]
	cc1 <- which(is.na(coal[i,])) 
		if (length(cc1)>0){
		for (k in 1:length(cc1)){
			rg <- which(tabsC[i2,]>0)
			for (m in 1:length(rg)){
				probCD[i1,which(locusiN[i1,]==cc1[k])][rg[m]] <- probCD[i1,which(locusiN[i1,]==cc1[k])][rg[m]] + C * tabsC[i2,rg[m]]}    
				probCD[i1,which(locusiN[i1,]==cc1[k])] [unique(corrCDi[i1,][which((locusiN[i1,]) == cc1[k])])] <- 0
				probCD[i2,which(locusiN[i2,]==cc1[k])] <- 0 }}}


## Coalescence (rate c)  
probCoalx <- integer(0)
for (m in 1:dim(coal)[1]){
	r1 <- combn(Sp, 2)[1,m]
	r2 <- combn(Sp, 2)[2,m]
	probCoal1 <- integer(0)
	c1 <- which(coal[m,]>0)
		for (i in c1){
		s1 <- unique(corrCDi[r1,which(locusiN[r1,]==coal[m,i])])
		s2 <- unique(corrCDi[(r2),which(locusiN[(r2),]==coal[m,i])])
			if (length(s1)>0 & length(s2)>0){
			if (s1==s2) { probCoal1 <- c(probCoal1,Tc)
					} else { probCoal1 <- c(probCoal1,0)}}}
probCoal <- integer(length(coal[1,]))
probCoal[which(coal[m,]>0)] <- probCoal1
probCoalx <- rbind(probCoalx,probCoal)}	

## Event sampling
probT1 <- integer(0)
for (i in 1:dim(probCD)[1]){
		probT1 <- c(probT1,probCD[i,])}
probT2 <- integer(0)
for (i in 1:dim(probCoalx)[1]){
		probT2 <- c(probT2,probCoalx[i,])}

probT <- c(probT1,probT2)
ss <- sample(1:length(probT),1,prob=probT)

## Time sampling

t1 <- -log(runif(1, min = 0, max = 1)) * 1/sum(probT)
times <- c(times, t1)

## Event consequences on gene locations
targ <- ceiling(ss/dim(probCD)[2])

if (targ<(Sp+1)){
ssx <- ss-(targ*dim(probCD)[2]-dim(probCD)[2])

# genome 1
	locS <- locusiN[targ,ssx]
	locusj1 <- locusiN[targ,1:(N*length(tabs[1,]))]
	corrCDi1_b <- corrCDi[targ,1:(N*length(tabs[1,]))]
	corrCDi1_b <- corrCDi1_b[-which(locusj1 == locusiN[targ,ssx])]
	locusj1 <- locusj1[-which(locusj1 == locusiN[targ,ssx])]
	rk <- which(corrCDi[targ,]==corrCDj[targ,ssx])[1]
			if (is.na(rk)) { 
				if (length(corrCDi1_b) == length(which(corrCDi1_b > corrCDj[targ,ssx]))) { locusj1 <- c(rep(as.numeric(locusiN[targ,ssx]),length(tabs[1,])),locusj1)}
				if (length(corrCDi1_b) == length(which(corrCDi1_b < corrCDj[targ,ssx]))) { locusj1 <- c(locusj1,rep(as.numeric(locusiN[targ,ssx]),length(tabs[1,])))}
	 			if (length(which(corrCDi1_b< corrCDj[targ,ssx]))>0 & length(which(corrCDi1_b> corrCDj[targ,ssx]))>0) {
					locusj1 <- c(locusj1[1:length(which(corrCDi1_b< corrCDj[targ,ssx]))],rep(as.numeric(locusiN[targ,ssx]),length(tabs[1,])),locusj1[(length(which(corrCDi1_b< corrCDj[targ,ssx]))+1):length(locusj1)])}
				 } else { locusj1 <- c(locusj1[0:(rk -1)],rep(as.numeric(locusiN[targ,ssx]),length(tabs[1,])),locusj1[rk:length(locusj1)])}

	locusj2 <- unique(locusj1[1:length(1:(N*length(tabs[1,])))])
	
	locusiN[targ,] <- c(locusj1[1:length(1:(N*length(tabs[1,])))],locusj2)
	tabs[targ,corrCDi[targ,ssx]] <- tabs[targ,corrCDi[targ,ssx]]-1
	tabs[targ,corrCDj[targ,ssx]] <- tabs[targ,corrCDj[targ,ssx]]+1
	coalT <- c(coalT,NA)
	coalPair <- rbind(coalPair,cbind(NA,NA))

# coalesced loci
		if (length(which(tabCoal == locS))>0) {
			cb1 <- which(combn(Sp, 2)[1,]==targ)
			cb2 <- which(combn(Sp, 2)[2,]==targ)
			cb <- c(cb1,cb2)
		
		if (length(cb)>0){
		for (i in 1:length(cb)){
			if (length(which(which(is.na(coal[cb[i],])==TRUE)==locS))>0){
				targ2 <- combn(Sp, 2)[2,cb[i]]
					if (targ2 == targ) {targ2 <- combn(Sp, 2)[1,cb[i]]}
				locusj1 <- locusiN[targ2,1:(N*length(tabs[1,]))]
				corrCDi2_b <- corrCDi[targ2,1:(N*length(tabs[1,]))]
				corrCDi2_b <- corrCDi2_b[-which(locusj1 == locS)]
				locusj1 <- locusj1[-which(locusj1 == locS)]
				rk <- which(corrCDi[targ2,]==corrCDj[targ,ssx])[1]
						if (is.na(rk)) { 
								if (length(corrCDi2_b) == length(which(corrCDi2_b > corrCDj[targ,ssx]))) { locusj1 <- c(rep(locS,length(tabs[1,])),locusj1)}
								if (length(corrCDi2_b) == length(which(corrCDi2_b < corrCDj[targ,ssx]))) { locusj1 <- c(locusj1,rep(locS,length(tabs[1,])))}
	 							if (length(which(corrCDi2_b< corrCDj[targ,ssx]))>0 & length(which(corrCDi2_b> corrCDj[targ,ssx]))>0) {
								locusj1 <- c(locusj1[1:length(which(corrCDi2_b< corrCDj[targ,ssx]))],rep(locS,length(tabs[1,])),locusj1[(length(which(corrCDi2_b< corrCDj[targ,ssx]))+1):length(locusj1)])}
						 } else { locusj1 <- c(locusj1[0:(rk -1)],rep(locS,length(tabs[1,])),locusj1[rk:length(locusj1)])}

				locusj2 <- unique(locusj1[1:length(1:(N*length(tabs[1,])))])
	
				locusiN[targ2,] <- c(locusj1[1:length(1:(N*length(tabs[1,])))],locusj2)
				tabs[targ2,corrCDi[targ,ssx]] <- tabs[targ2,corrCDi[targ,ssx]]-1  
				tabs[targ2,corrCDj[targ,ssx]] <- tabs[targ2,corrCDj[targ,ssx]]+1}}}}
			}

# if the event is a coalescence
if (targ>(Sp)){
ssC <- (ss - (dim(probCD)[2]*Sp))
ssC2 <- ceiling(ssC/N)
ssC3 <- ssC-(N*(ssC2-1))
	coal[ssC2,ssC3] <- NA
	tabCoal <- c(tabCoal,ssC3)
	coalT <- c(coalT,ssC3)
	coalPair <- rbind(coalPair,cbind(combn(Sp,2)[1,ssC2],combn(Sp,2)[2,ssC2]))}

for(y in 1:5){
for (i in 1:dim(coal)[2]){
	cl1 <- which(is.na(coal[,i])==TRUE)
		for(v in 1:length(cl1)){
			cla <- as.vector(combn(Sp, 2)[,cl1[v]])
			for (w in 1:length(cl1)){
				clb <- as.vector(combn(Sp, 2)[,cl1[w]])
				if (length(intersect(cla,clb))>0){
					cl2 <- unique(c(cla,clb))
					for(m in 1:dim(coal)[1]){
						m1 <- match(combn(Sp,2)[,m],cl2)
						m1 <- m1[!is.na(m1)]
						if (length(m1)==2) {coal[m,i] <- NA }}}}}}}

## Summary tables 
emptySp <- which(tabs[1,]==0)
for (i in 2:dim(tabs)[1]) {
		emptySp <- intersect(emptySp,which(tabs[i,]==0))}

		}

result <- list(coalescenceTab=coal,coalescedSpecies=coalPair,coalescedLocus=coalT,times=times)
return(result)

	}
```
