## Supplementary material for "The genomic view of diversification": GBD code: gbd.forward.R

```
gbd.f <- function(SpNb, LociNb, alpha, beta, delta, t) {

n <- LociNb
Sp <- SpNb

m <- alpha # genetic differentiation
r <- delta # gene flow
b <- beta # colonization

# all combination loci (numbers) * populations (letters)
p0 <- paste(1:n,rep((as.vector(sapply(letters,function(x) sapply(letters, function(y) paste0(x,y))))[1:Sp]),each=n), sep="")

# labels of genomes, at the beginning of the simulation only the first one exists
tabSp0 <- sapply(LETTERS,function(x) sapply(LETTERS, function(y) paste0(x,y)))[1:Sp]
tabSp <- tabSp0[1]

# matrix to record possible events depending on the existing populations
tab0 <- matrix(rep(0,(n*Sp*n*Sp)), ncol=n*Sp, nrow=n*Sp, byrow=TRUE)
diag(tab0) [1:n] <- 1 # at the beginning only one population (the AA population) exists
colnames(tab0) <- p0
rownames(tab0) <- p0
tab <- tab0

# table of existing loci
tabL <- cbind(rep(1:n, Sp),rep((as.vector(sapply(LETTERS,function(x) sapply(LETTERS, function(y) paste0(x,y))))[1:Sp]),each=n),rep("NA",(n*Sp)))
tabL [1:n,3] <- 0
rownames(tabL) <- paste(1:n,rep((sapply(letters,function(x) sapply(letters, function(y) paste0(x,y))))[1:Sp],each=n),sep="")

# table of co-adapted alleles
tabAssoc <- rbind(rep(0,2))

# table to record the score of co-adaptation between each gene lineage and each population
tabS <- matrix(rep(0,(Sp*n*Sp)), ncol=Sp, nrow=Sp*n)
tabS[1:n] <- n
colnames(tabS) <- (sapply(letters,function(x) sapply(letters, function(y) paste0(x,y))))[1:Sp]
rownames(tabS) <- paste(1:n,rep((sapply(letters,function(x) sapply(letters, function(y) paste0(x,y))))[1:Sp],each=n),sep="")

# result table
tabRes <- integer(0)

for (j in 1:t) {
	
# genetic differentiation
m2 <- which(!tabL[,3] == "NA")
probM <- cbind(rep(0,(n*Sp)))
probM[m2] <- m
rownames(probM) <- paste(1:n,rep(letters[1:Sp],each=n),sep="")

# gene flow
probR <- tab
diag(probR) <- 0

v1 <- which(probR==1)
for (i in v1){
	p1 <- i%%(n*Sp) # row
	p2 <- ceiling(i/(n*Sp)) # col
		if (p1==0){p1 <- n*Sp}
		if (p2==0){p2 <- n*Sp}

	probR[p1,p2] <- r * tabS[p1,ceiling(p2/n)]}

# colonization
probS <- rep(b, length(tabSp))

## event sampling
Prob <- c(as.numeric(probM),as.numeric(probR),as.numeric(probS))
s1 <- sample(c(1:length(Prob)), 1, prob=Prob)

## time sampling
times <- -log(runif(1, min = 0, max = 1)) * 1/sum(Prob)

## Event consequences
# genetic differentiation
if (s1 <= length(as.numeric(probM))) {        
	g2 <- tabL[,dim(tabL)[2]]
	g2[s1] <- j
	g2 <- g2[which(tabL[,2] == tabL[s1,2])]
	a2 <- unique(g2)
		for (i in 1:length(a2)){
			tabAssoc <- rbind(tabAssoc,sort(c(a2[i],j)))}

	A2 <- tabL[,dim(tabL)[2]]
	A2[s1] <- j
	tabL[,3] <- A2
	tabE <- c(p0[s1],j,"m")}

# gene flow
if (s1 > length(as.numeric(probM)) & s1 <= length(c(as.numeric(probM),as.numeric(probR)))) {    
	p1 <- ceiling((s1-length(as.numeric(probM)))/(n*Sp)) # col
	p2 <- (s1-length(as.numeric(probM)))%%(n*Sp) # row
		if (p1==0){p1 <- n*Sp}
		if (p2==0){p2 <- n*Sp}
	tabE <- c(p0[p1],p0[p2],"r")

	A2 <- tabL[,dim(tabL)[2]]
	A2 [p1] <- A2 [p2]
	tabL[,3] <- A2}

# colonization
if (s1 > length(c(as.numeric(probM),as.numeric(probR)))) {    
	tabSp <- c(tabSp,tabSp0[(length(tabSp)+1)])
	q1 <- s1-length(c(as.numeric(probM),as.numeric(probR)))
	genome1 <- which(tabL[,2] == tabSp[q1])
	genome2 <- which(tabL[,2] == tabSp[length(tabSp)])

	tabL[genome2 ,3] <- tabL[genome1 ,3]

	tabE <- c(tabSp[q1], tabSp[length(tabSp)], "s")

	for (i in 1:n) {
		tab [genome1[i],genome2[i]] <- 1
		tab [genome2[i],genome1[i]] <- 1}}

## Table (score of co-adaptation) update
# genetic differentiation
if (s1 <= length(as.numeric(probM))) { 
# rows
for (i in which(tabL[,2] == tabL[s1,2])) {
	for (k in 1:Sp) {

	g0 <- tabL[which(tabL[,2]==unique(tabL[,2])[k]),dim(tabL)[2]]
	m1 <- integer(0)
	for (q in 1:n) {
		for (p in 1:dim(tabAssoc)[1]) {
			compat1 <- c(match(tabAssoc[p,][1], sort(c(g0[q],tabL[i,dim(tabL)[2]]))[1]) , match(tabAssoc[p,][2], sort(c(g0[q],tabL[i,dim(tabL)[2]]))[2]))   
			if (length(which( compat1 == 1))==2) {m1 <- c(m1,1)} }}

tabS[i,k] <- length(m1)}}
	
# columns
for (i in 1:(n*Sp)) {
	for (k in ceiling(s1/n)) {

	g0 <- tabL[which(tabL[,2]==unique(tabL[,2])[k]),dim(tabL)[2]]
	m1 <- integer(0)
	for (q in 1:n) {
		for (p in 1:dim(tabAssoc)[1]) {
			compat1 <- c(match(tabAssoc[p,][1], sort(c(g0[q],tabL[i,dim(tabL)[2]]))[1]) , match(tabAssoc[p,][2], sort(c(g0[q],tabL[i,dim(tabL)[2]]))[2]))   
			if (length(which( compat1 == 1))==2) {m1 <- c(m1,1)} }}

tabS[i,k] <- length(m1)}}
	}

# gene flow
if (s1 > length(as.numeric(probM)) & s1 <= length(c(as.numeric(probM),as.numeric(probR)))) {    
# columns
p1 <- ceiling((s1-length(as.numeric(probM)))/(n*Sp)) # col
for (i in which(tabL[,2] == tabL[p1,2])) {
	for (k in ceiling(p1/n)) {

	g0 <- tabL[which(tabL[,2]==unique(tabL[,2])[k]),dim(tabL)[2]]
	m1 <- integer(0)
	for (q in 1:n) {
		for (p in 1:dim(tabAssoc)[1]) {
			compat1 <- c(match(tabAssoc[p,][1], sort(c(g0[q],tabL[i,dim(tabL)[2]]))[1]) , match(tabAssoc[p,][2], sort(c(g0[q],tabL[i,dim(tabL)[2]]))[2]))   
			if (length(which( compat1 == 1))==2) {m1 <- c(m1,1)} }}

tabS[i,k] <- length(m1)}}

# rows
for (i in p1) {
	for (k in 1:Sp) {

	g0 <- tabL[which(tabL[,2]==unique(tabL[,2])[k]),dim(tabL)[2]]
	m1 <- integer(0)
	for (q in 1:n) {
		for (p in 1:dim(tabAssoc)[1]) {
			compat1 <- c(match(tabAssoc[p,][1], sort(c(g0[q],tabL[i,dim(tabL)[2]]))[1]) , match(tabAssoc[p,][2], sort(c(g0[q],tabL[i,dim(tabL)[2]]))[2]))   
			if (length(which( compat1 == 1))==2) {m1 <- c(m1,1)} }}

tabS[i,k] <- length(m1)}}
	}

# colonization
if (s1 > length(c(as.numeric(probM),as.numeric(probR)))) {    #rows
	for (i in c(genome2)) {
	for (k in 1:Sp) {

	g0 <- tabL[which(tabL[,2]==unique(tabL[,2])[k]),dim(tabL)[2]]
	m1 <- integer(0)
	for (q in 1:n) {
		for (p in 1:dim(tabAssoc)[1]) {
			compat1 <- c(match(tabAssoc[p,][1], sort(c(g0[q],tabL[i,dim(tabL)[2]]))[1]) , match(tabAssoc[p,][2], sort(c(g0[q],tabL[i,dim(tabL)[2]]))[2]))   
			if (length(which( compat1 == 1))==2) {m1 <- c(m1,1)} }}

tabS[i,k] <- length(m1)}}
	}

if (s1 > length(c(as.numeric(probM),as.numeric(probR)))) {    #cols
	for (i in 1:(n*Sp)) {
	for (k in ceiling(genome2[1]/n)) {

	g0 <- tabL[which(tabL[,2]==unique(tabL[,2])[k]),dim(tabL)[2]]
	m1 <- integer(0)
	for (q in 1:n) {
		for (p in 1:dim(tabAssoc)[1]) {
			compat1 <- c(match(tabAssoc[p,][1], sort(c(g0[q],tabL[i,dim(tabL)[2]]))[1]) , match(tabAssoc[p,][2], sort(c(g0[q],tabL[i,dim(tabL)[2]]))[2]))   
			if (length(which( compat1 == 1))==2) {m1 <- c(m1,1)} }}

tabS[i,k] <- length(m1) 
			}
		}
	}

tabRes <- rbind(tabRes, matrix(c(tabE,times), ncol=4))

		}
		
return(tabRes)
		
	}
```
