## Supplementary material for "The genomic view of diversification": GBD code: make.tree.f.R

```
make.tree.f <- function(CoalSpTab, CoalLocusTab, CoalTime, SpNb, LociNb) {

coalPair <- CoalSpTab
coalL <- CoalLocusTab
coalT <- CoalTime
SpNb <- SpNb
LociNb <- LociNb

ag <- coalT[max(which(coalL>0))]

tree1 <- sim.bd.taxa.age(SpNb, 1, SpNb/ag, 0, age=ag)  
tp <- length(tree1[[1]]$tip.label)
nd <- tp-1

trees <- integer(0)
pair <- coalPair[which(coalL==1),]
tm <- coalT[which(coalL==1)]
nodes <- (nd+tp):(tp+1)

c2 <- integer()
for (i in 1:length(pair[,1])){
	c1 <- pair[i,]
		if (length(c1)>2){
			c1 <- c1[-3]
			c1[2] <- 10}
	c2 <- cbind(c2,c1)}	

c3b <- c2
c3b <- rbind(c3b,nodes)
for (i in 2:dim(c2)[2]){
for (j in 1:dim(c2)[1]){
	v1 <- c2[j,i]
	v2 <- c(match(v1,c2[1,1:(i-1)]),match(v1,c2[2,1:(i-1)]))
	v2 <- v2[!is.na(v2)]
		v2b <- c(match(nodes[v2],c3b[1,]),match(nodes[v2],c3b[2,]))
		v2b <- v2b[!is.na(v2b)]
		v3b <- v2b
		while (length(v2b)>0){ 
			v2b <- c(match(nodes[v2b],c3b[1,]),match(nodes[v2b],c3b[2,]))
			v2b <- v2b[!is.na(v2b)]
			v3b <- c(v3b,v2b)}
	if (length(v3b)>0) {v3 <- v3b[length(v3b)]} else {v3 <- v2}
				if (!is.na(v2[1])) {c3b[j,i] <- nodes[v3]}}}


c4 <- integer(0)
for (i in 1:dim(c2)[2]){
		c4 <- rbind(c4,rbind(cbind(c3b[3,i],c3b[1,i]),cbind(c3b[3,i],c3b[2,i])))}

c5 <- integer(0)
for (i in 1:dim(c4)[1]){
	t1 <- match(nodes,c4[i,])
	t1 <- t1[!is.na(t1)]
		if (length(t1)==1){
			t2 <- which(nodes==c4[i,1])
			c5 <- c(c5,tm[t2])}
		if (length(t1)>1) {
			t2 <- which(nodes==c4[i,1])
			t3 <- which(nodes==c4[i,2])
			c5 <- c(c5,tm[t2]-tm[t3])}}

tree1[[1]]$edge <- c4
tree1[[1]]$edge.length <- c5
tree1[[1]]$tip.label <- paste("t",1:tp,sep="")
plot(tree1[[1]])

return (tree1) }


## prepare files and run make.tree.f
make.tree.forward <- function (ResForward, SpNb, LociNb, SampledSp)  {

tabE0 <- ResForward
Sp <- SpNb
n <- LociNb
sampled.sp <- SampledSp
trees.f <- integer(0)

loci <- paste(1:n,rep((as.vector(sapply(letters,function(x) sapply(letters, function(y) paste0(x,y))))[1:Sp]),each=n), sep="")
species <- rep((as.vector(sapply(letters,function(x) sapply(letters, function(y) paste0(x,y))))[1:Sp]))
SPECIES <- rep((as.vector(sapply(LETTERS,function(x) sapply(LETTERS, function(y) paste0(x,y))))[1:Sp]))

s0 <- species[sampled.sp]
Sppairs <- combn(s0,2)
S0 <- SPECIES[sampled.sp]
SPpairs <- combn(S0,2)

if (length(which(is.na(tabE0) == TRUE)) == 0) {maxT <- dim(tabE0)[1]
	} else {maxT <- min(c(which(is.na(tabE0[,2])==TRUE)),which(is.na(tabE0[,3])==TRUE)) }

tabE0 <- tabE0[1:maxT,]

r1 <- cbind(loci,rep((as.vector(sapply(LETTERS,function(x) sapply(LETTERS, function(y) paste0(x,y))))[1:Sp]),each=n))
t <-  dim(tabE0)[1]

## In order to trace back the path of gene lineages, colonization events are transformed as gene flow for each gene lineage. 
events <- integer(0)
for (k in 1:length(sampled.sp)){
g0 <- loci [seq(1,(n*Sp),n)[sampled.sp[k]]:(seq(1,(n*Sp),n)[sampled.sp[k]]+(n-1))]
	
	for(i in 1:length(g0)) {
	g <- g0[i]
	sg <- g
	tg <- dim(tabE0)[1]

	tabE2 <- tabE0
	rp <- which(tabE2[,3]=="s")
		
		for(j in 1:length(rp)){
			rp1 <- r1[which(r1[,2] == tabE2[rp[j],2]) [i],1]
			rp2 <- r1[which(r1[,2] == tabE2[rp[j],1]) [i],1]
			tabE2[rp[j],c(1,2,3)] <- c(rp1,rp2,"r")}

			Eg <- intersect(which(tabE2[,1] == g), which (tabE2[1:t,3] == "r"))
				if(length(Eg) > 0){
					while (length(which(Eg< tg[length(tg)]))>0) {
						Eg2 <- Eg[which(Eg<tg[length(tg)])]
      					sg <- c(sg,as.character(tabE2[Eg2[length(Eg2)],2]))
						tg <- c(tg,Eg2[length(Eg2)])
						g <- sg[length(sg)]
						Eg <- intersect(which(tabE2[,1] == g[1]), which (tabE2[1:t,3] == "r")) }}

		events <- c(events,tg) }}

events <- unique(events)

## Coalescence time selection and production of imput tables for the function "make.tree.R".
T1 <- integer(0)
for (j in 1:n){
locus0 <- j
coals <- integer(0)
coalsTab <- integer(0)
	for (i in 1:dim(Sppairs)[2]){
		c1 <- paste(locus0,Sppairs[1,i],sep="")
		c2 <- paste(locus0,Sppairs[2,i],sep="")
		C1 <- SPpairs[1,i]
		C2 <- SPpairs[2,i]
		coals <- c(coals,max(c(intersect(which(tabE0 [,1] == c1),which(tabE0 [,2] == c2)),intersect(which(tabE0 [,2] == c1),which(tabE0 [,1] == c2)),max(c(which(tabE0 [,2] == C1),which(tabE0 [,2] == C2)))  )))}
	coals [which(coals == "-Inf")] <- NA

P1 <- integer(0)
P2 <- integer(0)
n1 <- integer(0)
nn <- 1
while(length(n1)<(length(sampled.sp)-1)) {
	n0 <- unique(match(sort(coals),coals))
	p1 <- Sppairs [1,n0[nn]]
	p2 <- Sppairs [2,n0[nn]]
	if (length(c(which(unique(P1) == p1),which(unique(P2) == p2),which(unique(P2) == p1),which(unique(P1) == p2)))<2) {
		n1 <- c(n1,n0[nn])
		P1 <- c(P1,p1)
		P2 <- c(P2,p2)}
	nn <- nn+1}

if (length(which(is.na(n1))) > 0) {
	n1 <- n1[-which(is.na(n1))] 
	n2 <- setdiff(n0,n1)
	L1 <- integer(0)
	nn <- 1
	for (y in 1:length(n2)) {
		p1 <- Sppairs [1,n2[nn]]
		p2 <- Sppairs [2,n2[nn]]
		L1 <- c(L1,length(c(which(P1 == p1),which(P1 == p2),which(P2 == p1),which(P2 == p2))))
		nn <- nn+1}
	n1 <- c(n1,n2[which(L1==max(L1))[1]])}

Coals <- rep(NA,length(coals))
Coals [n1] <- coals[n1]
coalsTab <- rbind(coalsTab,cbind(Coals,rep(locus0, length(coals)), Sppairs[1,], Sppairs[2,]))

coalsTab[,3] <- match(coalsTab [,3],species)
coalsTab[,4] <- match(coalsTab [,4],species)

# Controls for some particular cases
if(length(unique(c(as.numeric(coalsTab[,3]),as.numeric(coalsTab[,4]))))<length(sampled.sp)){
	v1 <- match(sort(coalsTab[,1]),coalsTab[,1])
	tv <- rbind(coalsTab[v1[1],3:4])
	for (i in 2:length(v1)) {
		tv0 <- coalsTab[v1[i],3:4]
			if (length(c(which (tv[,1]==tv0[1]), which (tv[,2]==tv0[1]))) > 0  & length(c(which (tv[,1]==tv0[2]), which (tv[,2]==tv0[2]))) > 0) {
			coalsTab[v1[i],1] <- NA} else {tv <- rbind(tv,coalsTab[v1[i],3:4]) }}
		
	miss <- which(is.na(match(c(1:length(sampled.sp)),(unique(as.numeric(coalsTab[,3],coalsTab[,4])))))==TRUE)
	missL <- c(which(coalsTab[,3] == miss),which(coalsTab[,4] == miss)) [min(which(coals[c(which(coalsTab[,3] == miss),which(coalsTab[,4] == miss))]>0))]
	coalsTab[missL,1] <- coals[missL]
		}

coalPair <- cbind(rep(NA, maxT),rep(NA, maxT))
coalLocus <- rep(NA, maxT)
ContinuousT <- rep(NA, maxT)
rg <- which(coalsTab[,1]>0)
for (i in 1:length(rg)) {
	coalPair [as.numeric(coalsTab[rg[i],1]),] <- as.numeric(coalsTab[rg[i],3:4])
	coalLocus [as.numeric(coalsTab[rg[i],1])] <- as.numeric(coalsTab[rg[i],2])
	ContinuousT <- cumsum(tabE0 [,4]) }
	
coalLocus [which(coalLocus == j)] <- 1
coalPair[,1] <- match(coalPair[,1], sampled.sp)
coalPair[,2] <- match(coalPair[,2], sampled.sp)

coalPair <- coalPair[sort(unique(c(events,which(coalLocus>0)))),]
ContinuousT <- as.numeric(ContinuousT[sort(unique(c(events,which(coalLocus>0))))])
coalLocus <- coalLocus[sort(unique(c(events,which(coalLocus>0))))]

trees.f <- c(trees.f, make.tree.f(coalPair, coalLocus, ContinuousT, length(sampled.sp) ,1)) 
	
		}
	return(trees.f)
	}
```
