## Supplementary material for "The genomic view of diversification": GBD code: read.me.r

```
# The genomic view of diversification (Julie Marin, Guillaume Achaz, Anton Crombach, Amaury Lambert)

## 1) The GBD-backward model
# The following parameters should be specified: SpNb (number of species), LociNb (number of loci), a (non-homologous attraction), b (homologous attraction), c (coalescence), d (erosion), and t (number of time units).
# The function will return 3 tables: 	
	# - the summary table of coalesced pair of species (row) and locus (columns) ($coalescenceTab), NA means that the locus i (columns) of the pair of species j (rows) have coalesced.
	# - the pair of species that have coalesced at each time unit ($coalescedSpecies), NA means that another event (not a coalescence) has occured at this time unit.
	# - the homologous locus that have coalesced at each time unit ($coalescedLocus), NA means that another event (not a coalescence) has occured at this time unit.

gbd.b (SpNb, LociNb, a, b, c, d, t) 

# Buiding trees with the outputs of gbd.b, the function will return a list containing all the gene (locus) trees.

make.tree (CoalSpTab, CoalLocusTab, SpNb, LociNb)

# example
library(TreePar)
library(phytools)
source("gbd.backward.R")
source("make.tree.R")

set.seed(1)
sim1 <- gbd.b(5, 10, 1, 1, 100, 2, 300) 
sim1$coalescenceTab # NAs indicate the loci that have coalesced, the simulation is complete if you have NAs in each cell, otherwise increase the parameter t.
gene.trees.simu1 <- make.tree(sim1$coalescedSpecies, sim1$coalescedLocus, sim1$times, 5, 10)

## 2) The GBD-forward model
# The following parameters should be specified: SpNb (number of species), LociNb (number of loci), alpha (genetic differentiation), beta (colonization), gamma (genetic drift), delta (gene flow) and t (number pf time units).
# The function will return a table indicating for each time unit the event that took place (3rd column, m = genetic differentiation, r = gene flow and s = colonization). If the event is a colonization, the replicated population is indicated in the first column. If the event if a mutation (genetic differentiation) the focal gene lineage and its population are indicated in the first column (for example "1aa" means that the locus 1 of in the population aa has undergone a mutation), the second column indicated the mutation number. If the event is an introgression (gene flow) the second column indicate the the gene lineage that is duplicated and that replaces the gene lineages indicated in the first column.

gbd.f(SpNb, LociNb, alpha, beta, delta, t) 

# Build trees
# The number of sampled species for which trees will be reconstruct has to be indicated. The label of species  can be specified (SampledSp <- c(1,5,3,8)) or chosen randomly (SampledSp <- sample (10, 4, replace=FALSE)).

make.tree.forward (ResForward, SpNb, LociNb, SampledSp) 

# example
library(TreePar)
library(phytools)
source("gbd.forward.R")
source("make.tree.f.R")

set.seed(1)
sim2 <- gbd.f(8, 4, 1, 1, 2, 300)
gene.trees.sim2 <- make.tree.forward(sim2, 8, 4, c(1:5)) 
```
